# Integration of synthetic microbial consortia based bioprocessing with pyrolysis for efficient conversion of cellulose to valuables

**DOI:** 10.1101/2022.06.26.497679

**Authors:** Chandrakant Joshi, Mahesh Kumar, Martyn Bennett, Jyotika Thakur, David J. Leak, Swati Sharma, Neil MacKinnon, Shyam Kumar Masakapalli

## Abstract

Improved technologies are needed for sustainable conversion of cellulosic waste to valuable products. Here we demonstrate the successful integration of a synthetic microbial consortium (SynCONS) based consolidated bioprocessing with pyrolysis to produce commodity chemicals from cellulose. Promising microbial partners were rationally identified from 7626 organisms via comparative metabolic mapping which led to establishing two promising SynCONS with abilities to convert cellulose to ethanol and lactate in bioreactors. The partners in the two SynCONS were a) the mesophilic fungus *Trichoderma reesei* grown sequentially with the thermophilic bacterium *Parageobacillus thermoglucosidasius* NCIMB 11955 (TrPt) and b) a thermophilic bacterium *Thermobifida fusca* grown together with *Parageobacillus thermoglucosidasius* NCIMB 11955 (TfPt). TrPt sequential bioprocessing resulted in 39% (g/g) cellulose consumption with product yields up to 9.3% g/g (ethanol + lactate). The TfPt co-cultures demonstrated a cellulose consumption of 30% (g/g) and combined yields of ethanol and lactic acid up to 23.7% g/g of consumed cellulose. The total product yields were further enhanced (51% g/g cellulose) when commercially available cellulases were used in place of *T. fusca*. Furthermore, when the metabolically engineered ethanol-producing strain of *P. thermoglucosidasius* TM242 (TfPt242) was substituted in the thermophilic TfPt co-culture consortium, ethanol yields were substantially higher (32.7% g/g of consumed cellulose). Finally, subjecting the residual cellulose and microbial biomass to pyrolysis resulted in carbon material with physicochemical properties similar to commercially available activated carbon as analysed using Scanning Electron Microscopy, X-Ray Diffraction and Raman spectroscopy. Overall, the integration of this synthetic microbial consortia-based bioprocessing strategy with pyrolysis demonstrated a promising strategy for conversion of waste cellulose to chemicals, biofuels, and industrial carbon potentially suitable for several industrial applications.

## Introduction

Due to our limited global reserves of fossil fuels, the increasing costs of oil extraction and the detrimental environmental effect of burning fossil fuels it is imperative to find alternative fuel and energy sources.^1,2^ Lignocellulosic biomass represents the most abundant renewable source of fixed carbon in the biosphere.^3,4^ It is an inexpensive and p more sustainable feedstock for producing useful bulk chemicals such as: bioethanol, biodiesel, lactic acid, iso-butanol, higher chain fatty acids etc.^3,5^ Indeed, large quantities of lignocellulosic waste are generated from agriculture, forest and industrial sources such as food processing and herbal product industries every year. The conversion of these renewable waste materials into useful chemicals provides a solution which addresses the challenges of sustainability.^6^ Bioprocessing of the cellulosic component of lignocellulose to added-value chemicals involves four major steps: 1) the physicochemical pre-treatment of the biomass, 2) the microbial production or introduction of saccharifying enzymes, 3) enzymatic hydrolysis of the biomass polysaccharides, and 4) the fermentation of the hydrolysed sugars to target products. All of these steps add costs to industrial scale production.^3,7^ Furthermore, physicochemical pre-treatment processes can release compounds which inhibit subsequent processing. As a consequence, these processes can require multiple washing, polishing and separation steps, adding avoidable costs.^8^ Hence, to reduce the number of processing steps, time and released toxins, a strategy to integrate enzyme production, saccharification and fermentation, called consolidated bioprocessing (CBP), has been proposed.^9^

Genetic engineering can serve as an efficient tool to convert a microbe into an organism capable of CBP, but engineering a single organism with multiple capabilities (including production of saccharifying enzymes, hexose and pentose sugar fermentation and resistance to the inhibitory products released from biomass pre-treatment) is a challenging task.^7,10^ In nature, microbes exist in diverse and intricate communities, where different microbes are specialised for different metabolic roles and niches.^7,11^ As an alternative to genetic engineering, synthetic microbial consortium (SynCONS) based CBP, inspired by nature, have also been explored.^10,12–14^

In a synthetic consortium for cellulose bioprocessing, two microbial species are selected with specialised functions, one dedicated to saccharification and the other to fermentation.^9^ Several successful studies have been reported using synthetic consortia based CBP in the last decade.^5,15–17^ The majority of the studies are focused on mesophilic consortia, with few on thermophilic consortia. Thermophilic bioprocesses involve naturally robust and thermostable enzymes beneficial for harsh industrial processing.^18–20^ Rapid fermentation is exothermic, but as thermophilic bioprocess runs at higher temperature this removes the need for cooling. Moreover, thermophilic fermentation can also aid in minimizing microbial contamination, and can assist downstream product recovery.^18–21^ Additionally, thermophilic microbes typically secrete a smaller set of enzymes than mesophilic fungi, for which they compensate by transporting and internally metabolising oligosaccharides.^22,23^ Owing to these advantages, cellulosic bioprocessing using thermophilic microbes has recently started to attract attention.

Notable thermophilic anaerobes can convert cellulose to fermentation products, however their natural yields are frequently sub-optimal from an industrial perspective. These lower yields can be primarily attributed to: (i) their nature of performing mixed acid fermentation where a main product is always accompanied with other products,^24–28^ and (ii) their sensitivity to increasing product concentrations, which in turn leads to fermentation inhibition.^24–30^ Consequently, many investigations have focused on developing genetically engineered thermophiles for improved bioprocessing.^31–34^ However, little progress has been made in improving these product yields in the context of thermophilic consortia, using either wild type or transgenic thermophiles, for cellulose bioprocessing.

In addition to desirable chemicals, the microbial bioprocessing of lignocellulosic materials also generates solid residues, typically composed of the remaining substrate and microbial biomass after bioprocessing. It is challenging to dispose of the large quantities of this residual slurry, given that disposal of this secondary waste may entail significant resources as well as raise issues of environmental pollution. While there are limited technologies to reuse the slurry via further anaerobic digestion processes, the associated challenges involve large volumes of water that ideally needs recycling. These factors pose challenges in large scale application of lignocellulosic bioprocessing.^35^. To address these challenges, we have integrated a pyrolysis technique, commonly used as a waste treatment strategy, for the complete utilisation of the residual biomass.

Pyrolysis is defined as the thermochemical decomposition of organic materials such as natural and synthetic polymers, gaseous hydrocarbons and some liquid petroleum by-products.^36^ This is carried out by heating the material in the absence of oxygen at temperatures typically above 500 °C.^37^ After pyrolysis, generally three types of products are generated: pyrolysis oils, flammable gases and solid carbon. The pyrolysis temperature and the gaseous environment in which the material is heated can be varied depending upon the application area. As a result, pyrolysis parameters can be tuned to generate high-quality carbon materials with potential commercial applications. For example, in the case of urban solid waste treatment, the commonly employed pyrolysis temperature is around 600 °C. On the other hand, for microfabrication purposes, the temperatures used are ≥ 900 °C and the environment is strictly inert.^37^ Notably, just by increasing the pyrolysis temperature and maintaining an inert environment, the quality of the resulting carbon can be drastically improved. Such a high-temperature inert pyrolysis is used for the commercial production of advanced carbon materials including: glass-like carbon, activated carbon and carbon fibers.^38^ Additionally, the small fraction of oils and gases produced during pyrolysis can be separately collected using any commercially available reactor in the case of very large biomass quantities. The resulting carbon materials can therefore potentially be used in multiple commercial applications based on their properties.

The integration of pyrolysis with microbial bioprocessing potentially increases the overall production of commercial valuables from this process. Moreover, it enables the safe use of transgenic microbes for lignocellulose degradation, since the high pyrolysis temperatures destroys the microbes, minimizing the need for safe disposal. While microbial bioprocessing and pyrolysis are two major techniques employed in parallel for biomass treatment, their integration by pyrolyzing the residuals at the end of bioprocess have not been reported. Conversely, some studies have been reported on the bioprocessing of products of biomass pyrolysis.^39–42^

In this work, we have integrated SynCONS based bioprocessing with pyrolysis for the complete utilisation of cellulose (Avicel) (Figure 1). We have successfully designed and established two synthetic microbial consortia (SynCONS) using the *in-silico* G2KO pipeline.^43^ Furthermore, we have optimised and validated two bioprocessing approaches for the conversion of cellulose to useful compounds. The first approach involves a sequential mesophile-thermophile:fungal-bacterial consolidated bioprocessing strategy (TrPt) combining the *Trichoderma reesei* QM6a and *Parageobacillus thermoglucosidasius* NCIMB 11955 for mixed acid fermentation of the resulting sugars. The second approach is involves a thermophile-thermophile:bacterial-bacterial consolidated bioprocessing strategy (TfPt) with cocultures of *Thermobifida fusca* ATCC 27730 and *Parageobacillus thermoglucosidasius* NCIMB 11955 for mixed fermentation of the resulting sugars. To further establish the efficiency of the designed and optimised consortia, we also utilised an engineered ethanol production strain of *Parageobacillus thermoglucosidasius* TM242 to perform ethanologenic fermentation in the TfPt strategy.^44^ The efficiency and yields of these synthetic consortia were compared with a bioprocess where commercially available cellulases were used to degrade cellulose before the sugars were fermented by *P. thermoglucosidasius* NCIMB 11955.

**Figure 1:**
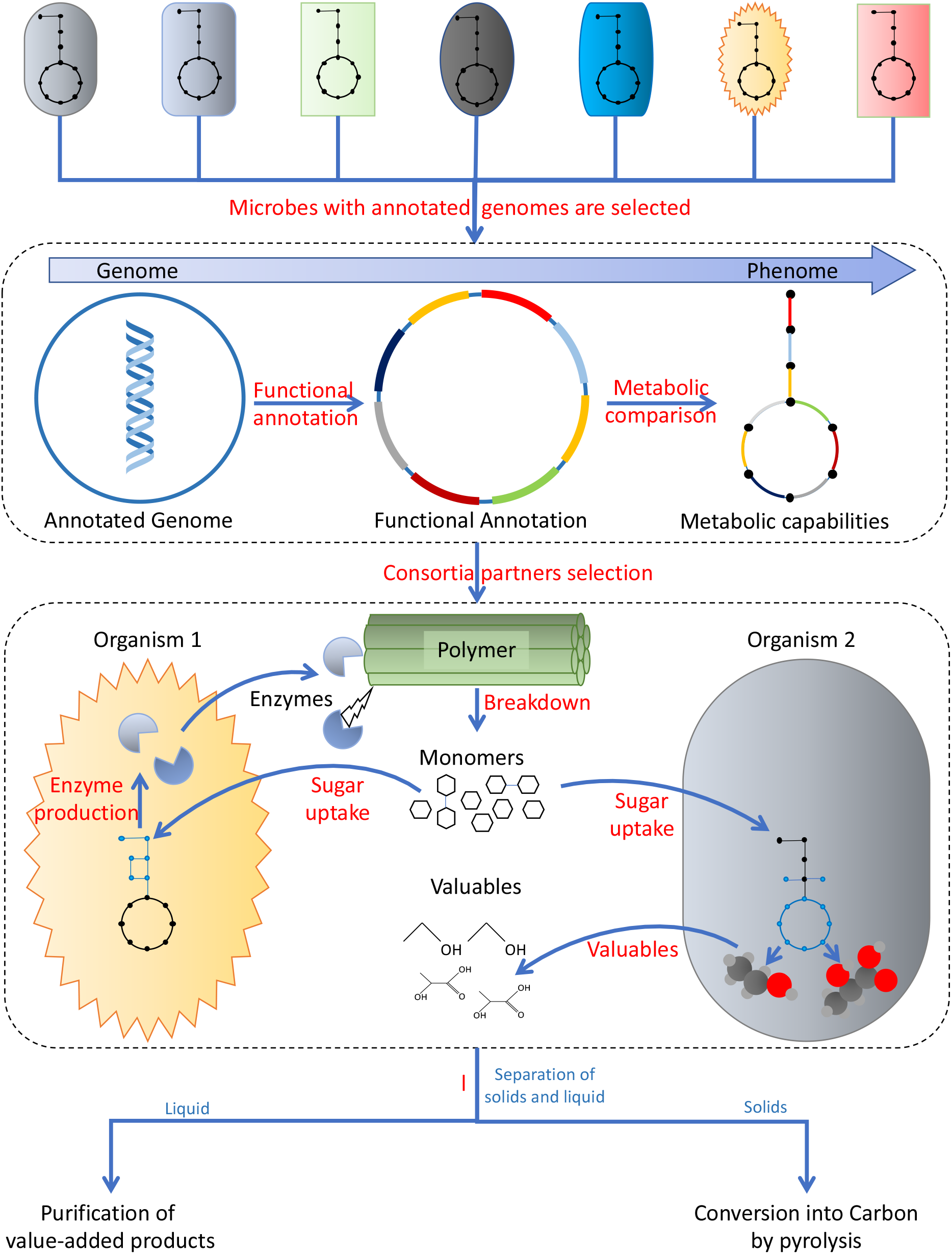
Rational strategy of integrating Synthetic microbial consortia (SynCONS) with pyrolysis. Microbes with functionally annotated genomes are selected from the literature based on desirable metabolic capabilities. Functional annotation of the genes allows visual comparison of the metabolic capabilities of different organisms using tools like G2KO and KEGG mapper. The best consortium microbe partners among the selected microbes are selected based on their metabolic capabilities and their physico-chemical compatibility. The consortia are then validated physically in bioprocessing experiments and further optimised.

Overall, a holistic approach of a “minimum-waste” conversion of cellulosic biomass was rationally established by integrating SynCONS based bioprocessing with pyrolysis, which resulted in an efficient conversion of waste cellulose to chemicals, biofuels, and industrial carbon suitable for various downstream applications.

## Experimental

### Designing SynCONS for cellulose to value-added chemicals

At the time of writing, there were 7626 genomes of a variety of organisms available on the KEGG database. Out of these, 6911 microbial genomes were selected, and finally narrowed to a list of 218 organisms by searching the literature for reported cellulose degradation and/or fermentation capabilities at the genus level. The metabolic capabilities of the selected 218 organisms were compared using the online pipeline G2KO and KEGG mapper. The visual comparisons from the KEGG mapper were manually quantified and a pathway completeness score was generated. Leading organisms demonstrating high completeness for known cellulose degradation and fermentation pathways were investigated further for their feasibility as potential consortium partners, based on their optimum growth and nutritional parameters.

### Microbial strains and media components

The microbes employed in this study were selected to perform one of two roles in the consortia, either cellulose degradation or sugar fermentation. The mesophilic filamentous fungus *T. reesei* QM6a and thermophilic bacteria *T. fusca* ATCC 27730 were used as cellulose degraders while the lactic acid producing thermophilic bacteria *P. thermoglucosidasius* NCIMB 11955 was used to ferment the resulting sugars. *T. reesei* QM6a was procured from the National Collection of Industrial Microorganisms (NCIM), National Chemical Laboratory (NCL), Pune, India. *P. thermoglucosidasius* strains NCIMB 11955 and TM242 were available in house. *P. thermoglucosidasius* TM242 was genetically engineered from wild type *P. thermoglucosidasius* NCIMB 11955 by Cripps et al. (2009) and contained deletions in the lactate dehydrogenase gene *ldhA* and pyruvate formate lyase gene, together with upregulated expression of the pyruvate dehydrogenase operon.^44^ *T. fusca* ATCC 27730 was obtained from Microbial Type Culture Collection and Gene Bank (MTCC), Institute of Microbial Technology (IMTECH), Chandigarh, India.

Several different microbial culture media were used in this study: potato dextrose agar (PDA) medium, Trichoderma minimal medium (TMM), ammonium salt medium (ASM), tryptone-yeast extract (2TY) liquid medium, 2TY agar medium, Hagerdahl medium and tryptone yeast glucose agar (TYG). Detailed compositions of each medium can be found in supplementary materials section 2. All the prepared media were autoclaved at 121 °C for 20 min and thermo-degradable components (biotin and thiamine) were filter sterilised through 0.22-micron filters. The *Trichoderma reesei* commercial cellulases were obtained from Sigma (C2730).

### Culturing, maintenance, and inoculum preparation

The *T. reesei* cells were grown and maintained on PDA media plates at 28 °C, *P. thermoglucosidasius* strains on 2TY agar plates at 60 °C and *T. fusca* on TYG plates at 55 °C, all in static incubators. Three-day plate-grown *T. reesei* cultures was used to inoculate liquid cultures for the sequential bioprocessing of cellulose. *T. reesei* fungal discs with a diameter of 9.5 mm were cut out from the PDA plate using a cork-borer. Agar was separated from the disc and the mycelial mat was used as inoculum. For bacterial culture, *P. thermoglucosidasius* strains were first grown from a plate in 15 mL of 2TY medium (50 mL culture tubes) for 24 hours at 55 °C and 180 RPM shaking, then centrifuged (ThermoFisher Scientific Heraeus Multifuge X3R with Fiberlite F14-6×250 LE rotor) at 2276 g for 5 min and the supernatant discarded. The resulting cell pellet was washed with ASM without any carbon source by resuspending the cells in 10 mL ASM (without carbon source) and centrifuging once at 2276 g for 5 min. This cell pellet was used as the inoculum for fermentation in the sequential microbial bioprocessing of cellulose. *T. fusca* inoculum was grown in Hagerdahl medium for 3 days at 55 °C and 180 RPM shaking with 2% glucose as a carbon source and subsequently used in experiments.

### Synthetic microbial consortia-based bioprocessing and its optimisation

For proving the concept of fungal-bacterial sequential bioprocessing to produce lactic acid, 10 mycelial discs of *T. reesei* were inoculated in TMM media (pH 4.8) with 2 % Avicel and incubated at 28 °C, 150 rpm for 5 days in 50 mL tubes with culture volume of 10 mL. After 5 days, the culture pH was adjusted to 6.8 using KOH and 10 mL 2X ASM media without glucose was added to support the growth of *P. thermoglucosidasius*. 1 mL *P. thermoglucosidasius* NCIMB 11955 inoculum (OD_600_, 0.211) was then added to the same tubes, and the culture was incubated at 55 °C for 48 hours at 180 RPM. This effectively killed the *T. reesei* but allowed *P. thermoglucosidasius* to ferment the resultant sugars. Finally, the culture was harvested after 48 h. The harvested culture was used to confirm lactic acid production.

Process conditions were subsequently refined in order to improve the final product yield. The fungal inoculum concentration was optimised for sugar release from cellulose by taking different numbers of mycelial discs ranging from 1 to 14 (Suppl. Fig 2a). The cultures were collected after 5 days of growth and used to quantify sugars released from cellulose enzymatic breakdown. The optimal growth time for the fungus to degrade cellulose was determined by growing the *T. reesei* with optimal fungal inoculum at 28 °C and 150 rpm for up to 10 days. The culture samples were collected each day and subjected to total reducing sugar analysis. After growing the fungus with optimised inoculum at 28 °C for the optimised time period, the pH was adjusted to approximately 4.8 and then additional saccharification with the *T. reesei* enzymes was carried out by incubating the culture at 50 °C, 150 rpm for 24 hours. In these 24 hours, the *T. reesei* cells cannot survive because of high temperature, but the enzymes produced by *T. reesei* work optimally at 50 °C and continue to breakdown cellulose. ^45^ Culture samples were collected every 6 hours during this additional time and total reducing sugar content was analysed. For bacterial inoculum optimization, a monoculture of *P. thermoglucosidasius* NCIMB 11955 was grown in ASM with 0.5 % glucose as carbon source. The pre-cultured *P. thermoglucosidasius* was used as inoculum as described above (in the section “Culturing, maintenance and inoculum preparation”). Inoculum (OD, 0.254) concentrations 10 %, 20 % and 30 % (v/v) of culture media (20 mL), were used for optimisation. Samples were collected after 24 hours and lactic acid concentration was measured. To obtain the highest product yield the 10 % v/v inoculum of *P. thermoglucosidasius* NCIMB 11955 was grown in ASM with 0.5 % glucose and samples were collected every 12 hours, up to48 hours after inoculation. Each sample was then subjected to HPLC analysis for lactic acid quantification. After this optimisation, sequential bioprocessing of cellulose was performed with the optimized conditions and lactic acid was quantified.

To optimise the *T. fusca* inoculum for maximum sugar release from cellulose, 10 mL of Hagerdahl medium (with cellulose as carbon source) was put in Corning minibioreactor tubes (50 mL tubes with a membrane in the cap to facilitate the air exchange) and inoculated with different concentrations (10 %,20 % and 30 %) of primary culture at 55 °C. Samples were taken every 24 hours, and the sugar content was determined using DNS assay.

### Bioprocessing of Avicel with optimised parameters

The fungal-bacterial (TrPt) sequential bioprocessing of cellulose in bioreactors (ApplikonMiniBio 250 mL) was done using the optimised parameters obtained as described above. In this sequential bioprocessing, there are two distinct phases; first the cellulose degradation phase where *T. reesei* breaks down the cellulose into glucose and cellobiose, and second where *P. thermoglucosidasius* is added to ferment those sugars into acetate, lactate, formate and ethanol. These two phases have different optimum parameters for microbial growth (Suppl. Fig. 1). In the cellulose degradation phase, agitator RPM was set to 200 for gentle mixing of the culture, allowing the fungal mycelia to grow without much disruption. The physicochemical parameters of: a 28 °C, pH 4.8, maximum air flow of 1 VVM (volume of gas per volume of liquid per minute) and a dissolved oxygen tension (DOT) of 20 % were set to allow optimum growth of the fungus. After 7 days the temperature was increased to 50 °C for 24 hours, which would inactivate the fungus but is optimum for fungal cellulase activity. For the second phase (fermentation), preheated 2X ASM media without a carbon source was added and parameters were changed to support *P. thermoglucosidasius* fermentation. Temperature was increased to 55 °C, pH was set to 6.8, and agitator RPM was increased to 300. Once conditions had equilibrated, *P. thermoglucosidasius* (10 % v/v) was added. After the *P. thermoglucosidasius* inoculation, aeration was set for 1 VVM for 4 hours, afterwards it was reduced to 0.02 VVM to support fermentation and cellulose degradation (Suppl. Table 1)._46–48_

For the bacterial-bacterial (TfPt) cellulose bioprocessing experiment in bioreactors (Applikon Minibio, 250 mL vessel), 80 mL of Hagerdahl minimal medium was added with 2 % w/v Avicel as a carbon source. The operational parameters (Suppl. Fig. 2) of 300 rpm, pH 7 and 55 °C were set. A DOT of 20 % was set with a maximum 1 VVM air flow allowed. Then 10 % v/v *T. fusca* inoculum was added. Triplicate samples of 1 mL were taken every 24 hours. At this point, preheated 2X ASM media (without carbon source) was added and *P. thermoglucosidasius* was inoculated (10 % v/v) into the bioreactor. After 4 hours of P. thermoglucosidasius inoculation in the bioreactors, aeration was reduced to 0.02 VVM to support fermentation and cellulose degradation (Suppl. Table 2).^46–48^

### Enzyme-bacterial Simultaneous Saccharification and Fermentation (SSF) bioprocessing of cellulose in the bioreactors

Optimisation of the enzyme loading for the simultaneous saccharification and fermentation was done in 50 mL Corning bioreactor tubes (4 biological replicates) at different cellulases concentrations of 7, 14, 21 and 28 units/mL with reaction times of 48 hours. Each tube (50 mL culture tubes) contained 10 mL citrate buffer (pH 4.8) and 2% w/v Avicel (Sigma). The samples were collected periodically and stored at -20 °C until further analysis. The DNS assay was performed to quantify the total sugars released (glucose equivalent) by the enzymatic breakdown of cellulose. The quantification analysis was done against the standard concentration of glucose.

SSF in the bioreactors was performed with optimum parameters of 7 unit/mL cellulase, ASM media with Avicel as sole carbon source, pH4.8 and 50 °C for 24 hours. After 24 hours the bioprocess parameters were adapted for the growth of *Parageobacillus thermoglucosidasius*. The temperature and pH were raised to 55 °C and 6.8 respectively, aeration was started and the *Parageobacillus thermoglucosidasius* was inoculated to the bioreactors. Aeration was 1 VVM for first 4 hours and 0.02 VVM afterwards throughout the bioprocessing, and DOT was set at 20%.

### Pyrolysis

After sequential bioprocessing of cellulose, the residual biomass was separated from the liquid by centrifugation (2276 g for 10 minutes) and dried in the oven at 200 °C for 4-5 hours. It was then pyrolysed in a tube furnace (JUPITER, India) under N_2_ flow (flowrate 0.8 L/min) at 900 °C with a temperature ramp rate of 5 °C per minute. Dwell time at the final 900 °C pyrolysis temperature was 1 hour.

### Estimation of inoculum amount, total biomass, total reducing sugar content and total sugar content

The initial inoculum and total residual biomass (cells and unutilised cellulose) of final cultures *T. reesei* bioprocessing were quantified gravimetrically.^7^ Estimation of total reducing sugar content was carried out by the DNS method as described by Minty *et al*. (2013).^7^ The culture supernatant (100 µL) was taken and diluted to 500 µL with DI water and the cellulose breakdown was quenched and analysed for reducing sugars by adding 1 mL of 3,5-dinitrosalicylic acid (DNS) reagent mix and incubating at 95 °C for 10 minutes.^49^ Absorbance was measured at 540 nm and a glucose standard was used to prepare the calibration curve. The total sugar content was estimated by the phenol-sulfuric assay. ^50^ The assay reaction mixture contained 30 µL of culture supernatant, 200 µL of concentrated sulfuric acid and 50 µL of 80 % aqueous phenol and the absorbance was recorded at 488 nm after a 30 minute incubation at 28 °C. A calibration curve was prepared using glucose as a standard and used for calculating the total sugar content.

### Total protein content

A Bio-Rad DC protein assay kit was used to perform total protein content in cellular and culture supernatants.^51^ The assay was carried out in a 96 well microtiter plate and was performed according to the manufacturer’s instructions. A calibration curve was prepared using a bovine serum albumin (BSA) standard of varying concentrations from 50-750 µg/mL. For the determination of extracellular protein content, 5 µL filtered culture sample was directly used as the sample for the protein assay. For the determination of cellular protein content, cell pellets were first washed thrice with DI water, then the pellet was frozen and crushed in liquid nitrogen, suspended in the extraction buffer (Tris-HCl/EDTA buffer, pH 8.0) and sonicated for 30 minutes to lyse the cells. After sonication, the solution mix was centrifuged at 14224 g (temperature, 4 °C) for 10 minutes. The resulting supernatant was used as the sample for cellular protein measurement.

### Enzyme assays

The activities of the endoglucanases, exoglucanases and β-glucosidases present in the commercial cellulase mixture were determined by the method of Minty *et al*. (2013).^7^ For the endoglucanase assay, the culture samples in 250 µL citrate buffer (50 mM, pH 4.8) were pre-incubated at 50 °C for 5 minutes. Carboxymethylcellulose (CMC) (1 % in 250 µL DI water) was added to initiate the reaction and the mixture was incubated for 30 minutes at 50 °C with continuous shaking at 900 rpm. Reactions were quenched with 1 mL of DNS reagent mix and then incubated at 95 °C for 10 minutes. The concentration of the resulting 3-amino, 5-nitrosalicylic acid (produced stoichiometrically from reducing sugars) was determined by measuring absorbance at 540 nm. Glucose was used as a calibration standard with 100 to 600 µM in an assay mix. One international unit (IU) of enzyme activity was defined as the amount of enzyme required to produce 1 µmol glucose-equivalent per minute. Exoglucanase activity was assayed in a similar way to the endoglucanase assay, except crystalline cellulose (Avicel® PH-101) was used as the substrate.^7^In the β-glucosidase assay, *p*-nitrophenyl-*β*-D-glucopyranoside (*p*NPG) was used as the substrate. The culture samples in 200 µL citrate buffer (50 mM, pH 4.8) were pre-incubated at 50 °C for 5 minutes, then the reaction was initiated by adding *p*NPG to 2.5 mM and the mixture incubated at 50 °C for 10 min. Finally, NaOH-glycine buffer (pH 10.8) was added to 0.2 M in the mixture to quench the reaction. The *p*-nitrophenol released in the assay was measured by its absorbance at 405 nm. A 0 to 0.25 mM *p*-nitrophenol calibration curve was prepared in the assay mix. One IU of β-glucosidase activity was defined as the amount of enzyme required to produce 1 µmol of p-nitrophenol per minute.

### HPLC analysis

Concentrations of lactic acid, glucose and ethanol were analysed by HPLC (Agilent Technology 1260 infinity) using a photodiode array (PDA) detector and refractive index detector (RID). Separation was achieved on a SUPELCOGEL™ C-610 column (9 µm, 30 cm x 7.8 mm) with 5 mM H_2_SO_4_ eluent. The HPLC was carried out with the following parameters: column temperature: 60 °C; flow rate: 0.6 mL/min; injection volume: 10 µl. For lactic acid detection and quantification, PDA detector was used at a detection wavelength of 210 nm and a run time of 40 minutes. Glucose and ethanol concentrations were determined using the RID, which was kept at 35 °C throughout the run. EZChrom Elite software was used for data analysis. The analyte was identified by comparing its retention time with that of pure standard as well as spiking samples with the standard compound. Quantification of the compound was performed on the basis of peak area through comparison with a calibration curve obtained with the respective standard.

### Proton (^1^H) NMR

Cell-free culture sample extracts obtained after sequential microbial bioprocessing of cellulose were also analysed by ^1^H NMR. In each case, 300 µL of filtered sample (filter mesh size: 0.2 µm) was diluted with 300 µL D_2_O and an internal standard of 100 µL D_2_O, containing 0.01 % w/v of 4,4-Dimethyl-4-silapentane-1-sulfonic acid (DSS) was. One-dimensional proton NMR was measured on a JEOL-ECX-500 MHz NMR Spectrometer. The measurement parameters were as follows: 45° pulse 5.37 μs, 5 s relaxation delay, 9.0 ppm spectral width, 1.45 s acquisition time, with 64 scans collected over a 16-minute data accumulation time. The residual water signal was suppressed with pre-saturation during the relaxation delay. The resulting spectra were phase and baseline corrected and then the chemical shift was calibrated using the DELTA 4.3.6 software.

### Raman spectroscopy, X-Ray Diffraction (XRD) analysis and Scanning Electron Microscopy (SEM)

Raman spectroscopy for the characterization of carbon (obtained by the pyrolysis of process residues) was carried out using a confocal Microscope Raman spectrometer (Horiba Scientific, XploRA ONE) with an excitation wavelength of 532 nm. XRD of carbon was carried out on a Rigaku SmartLab X-Ray Diffractometer equipped with a 9kW rotating anode X-Ray generator. SEM imaging was carried out on a JFEI Nova Nano SEM-450 Field Emission SEM system.

## Results

### Synthetic microbial consortia (SynCONS) with metabolic capabilities of converting cellulose to fermentative products established

The 218 organisms initially identified from the KEGG database were compared based on target metabolic capabilities and the complete results are presented in Suppl. Figure 9, and Suppl. Table S3. Two potential consortia with the metabolic ability of cellulose degradation and fermentation, either as cocultures or sequentially, were proposed from this analysis. Leading organisms with maximum cellulose degradation and fermentation pathways completeness are presented in Tables 1 and 2 respectively.

**Table 1:** Leading organisms for cellulose degradation pathway completeness, as highlighted in our *in-silico* study of 218 microbial genomes (See Suppl. Table S3).

| KEGG Code | Organism | Cellulose Degradation Completeness (%) | Optimum Temp. | Cellulases system/ growth | Optimum pH |
| --- | --- | --- | --- | --- | --- |
| tfu | *Thermobifida fusca*<sup>52</sup> | 86 | 50-55 °C | Aerobic | 7 |
| ccb | *Clostridium cellulovorans*<sup>53</sup> | 86 | 37 °C | Anaerobic | 7 |
| cac | *Clostridium acetobutylicum*<br>ATCC 824<sup>54</sup> | 86 | 35 °C | Anaerobic | 6.6 |
| cae | *Clostridium acetobutylicum*<br>DSM 1731<sup>55</sup> | 86 | 37 °C | Anaerobic | 7.1 |
| cay | *Clostridium acetobutylicum*<br>EA 2018<sup>56</sup> | 86 | 37 °C | Anaerobic | 5.0 |
| clb | *Clostridium sp.* BNL1100<sup>57</sup> | 86 | Mesophile | Anaerobic | NA |
| csr | *Clostridium*<br>*saccharoperbutylacetonicum*<br>N1-4<sup>58</sup> | 86 | 25 - 35 °C | Anaerobic | 5.6 – 6.7 |
| csb | *Clostridium*<br>*saccharobutylicum* DSM<br>13864<sup>58</sup> | 86 | 30 – 34 °C | Anaerobic | 6.2 - 7 |
| gjf | *Geobacillus genomosp. 3*<br>(*Geobacillus sp.* JF8)<sup>59</sup> | 86 | 60 °C | NA | NA |
| ttm | *Thermoanaerobacterium*<br>*thermosaccharolyticum* DSM<br>571<sup>60</sup> | 86 | 50 °C | Anaerobic | 6.8 |
| tsh | *Thermoanaerobacterium*<br>*saccharolyticum* JW/SL-YS485<sup>61</sup> | 86 | 55 °C | Anaerobic | 6.1 |
| acel | *Anaerocolumna cellulolytica*<br>SN021<sup>62</sup> | 86 | 35 °C | Anaerobic | 8.5 |
| cceu | *Cellulosimicrobium cellulans*<br>PSBB019<sup>63</sup> | 86 | Mesophile | Anaerobic | NA |
| bco | *Bacillus cellulolyticus* DSM<br>2522<sup>64</sup> | 86 | 37 °C | Aerobic | 9 - 10 |
| tre | *Trichoderma reesei* QM6a<sup>65</sup> | 71 | 28 °C | Aerobic | 4.8 |

**Table 2:** Leading organisms for fermentation pathway completeness, as highlighted in our *in-silico* study of 218 microbial genomes (See Suppl. Table S3).

| KEGG Code | Organism | Fermentation Completeness (%) | Optimum Temp. | Optimum pH |
| --- | --- | --- | --- | --- |
| cceu | *Cellulosimicrobium cellulans*<br>PSBB019<sup>63</sup> | 86 | Mesophile | NA |
| cga | *Cellulomonas gilvus* ATCC 13127<sup>66</sup> | 86 | 30 °C | NA |
| celh | *Cellulomonas sp.* H30R-01 | 86 | NA | NA |
| kll | *Klebsiella sp.* LTGPAF-6F | 86 | NA | NA |
| klw | *Klebsiella huaxiensis*<sup>67</sup> | 86 | 35 °C | NA |
| kok | *Klebsiella oxytoca* KONIH1 | 86 | NA | NA |
| koc | *Klebsiella oxytoca*CAV1374<sup>68</sup> | 86 | 30 °C | 3-9 |
| xyl | *Xylanimonas allomyrinae* 2JSPR-7<sup>69</sup> | 86 | 28–30 °C | 7.0–8.0 |
| cet | *Cellulosimicrobium sp.* TH-20<sup>70</sup> | 86 | 30 °C | 7 |
| gjf | *Geobacillus genomosp. 3*<br>(*Geobacillus* sp. JF8)<sup>59</sup> | 79 | 60 °C | NA |
| gth | *Parageobacillus*<br>*thermoglucosidasius* C56-YS93<sup>71</sup> | 71 | 75 °C | 6.4 |

### Fungal-bacterial sequential SynCONS of *Trichoderma reesei - Parageobacillus thermoglucosidasius* (TfPt) for cellulose bioprocessing

For the sequential TfPt SynCONS focused on lactic acid production as a proof of concept, the mesophilic fungus *T. reesei* NCIM942 was used as the cellulose degrader and the thermophilic bacterium *P. thermoglucosidasius* NCIMB 11955 was used as a sugar fermenter (Figure 2). Yields of lactic acid from this process were quantified by HPLC, GC-MS and proton NMR (Suppl. Fig. 4). To maximise the lactic acid yield, optimisation of the sequential bioprocessing was performed. The optimum fungal inoculum was found to be 10 mycelial discs (18.12 ± 0.2 mg dry weight) as this released the maximum amount of reducing sugar (Suppl. Fig. 5a). The optimal incubation time at 28 °C for maximum reducing sugar release was 7 days (168 h) (Suppl. Fig. 5b). The culture was further incubated at 50 °C (as media contained active endoglucanase, exoglucanase and *β*-glucosidase with activities 115.99 ± 5.82 U, 78.88 ± 3.94 U, and 23.94 ± 2.11 U, respectively) (Suppl. Fig. 6) and samples were taken periodically as detailed in the materials and methods. The results demonstrated the maximum total reducing sugar released through subsequent activity of *T. reesei* cellulases after 18 hours of incubation at 50 °C (Suppl. Fig. 5c). For fermentation, the optimised bacterial inoculum and incubation time was found to be 10 % (v/v of culture media) of OD_600_ 0.254 at 48 h, respectively (Suppl. Fig. 7a, b). The highest lactic acid titre obtained from the TrPt sequential bioprocessing under these conditions was 0.262 (± 0.015) g/L. The amount of generated biomass or solid residues after bioprocessing under these conditions was 121.84 (± 10.08) mg from 10 mL culture, which was subjected to pyrolysis.

**Figure 2:**
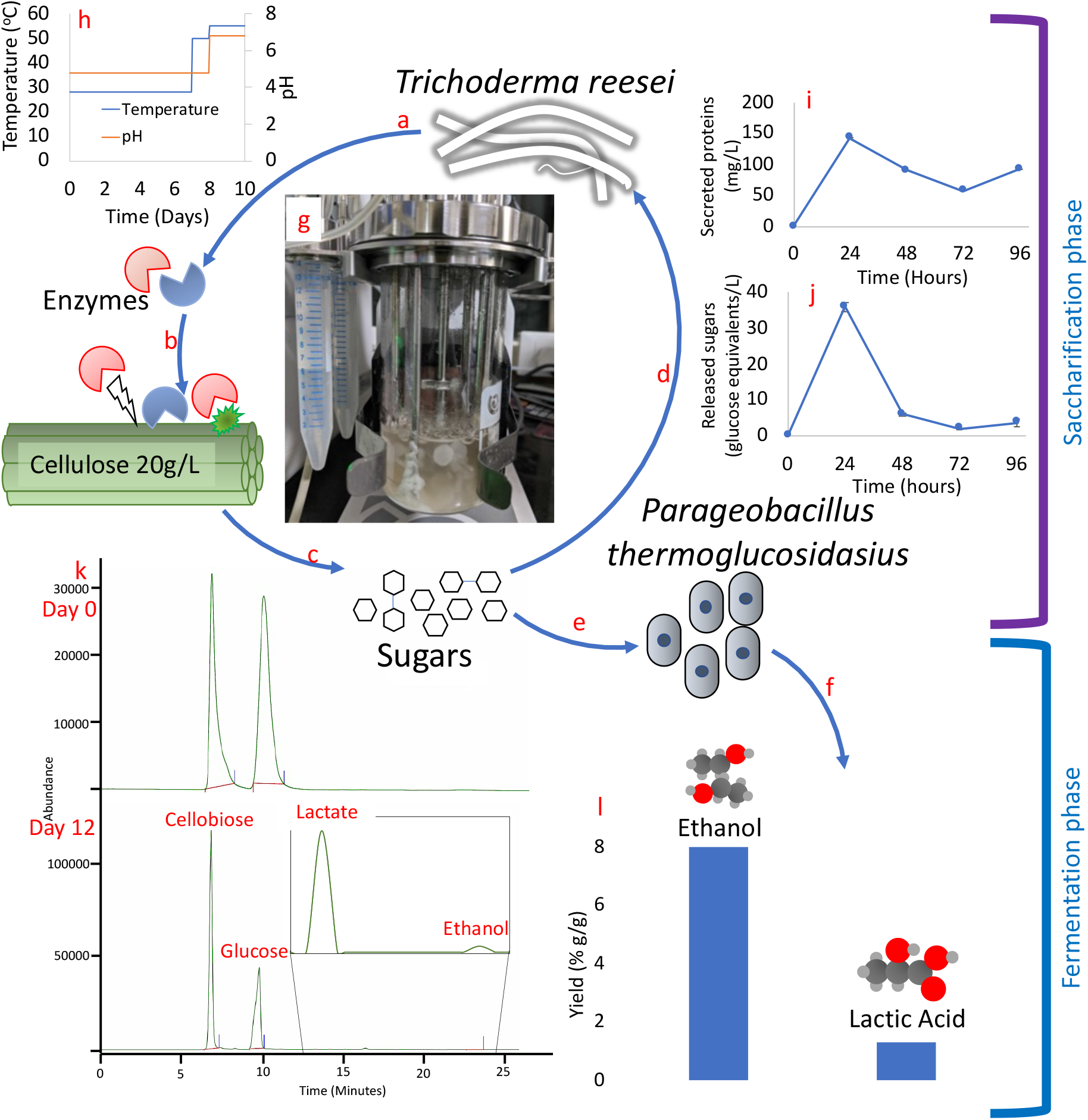
The SynCONS of the *Trichoderma reesei -Parageobacillus thermoglucosidasius* (TfPt) fungal-bacterial sequential microbial consortia for cellulose bioprocessing. The sequential consortium has two distinct phases: the saccharification phase (*T. reesei*) and the fermentation phase (*P. thermoglucosidasius*). a) *T. reesei* produces and secretes the cellulase enzymes, b) these enzymes break down the cellulose into simpler sugars, c) the simpler sugars (e.g., glucose and cellobiose) are released from cellulose breakdown, d) the sugars are taken up by *T. reesei* for growth, metabolism and enzyme production, e) after 7 days, *P. thermoglucosidasius* is introduced. It takes up the sugar, f) the sugars are being utilised by *P. thermoglucosidasius* for the biochemical production of ethanol and lactic acid focused on in this experiment. g) the bioreactor containing the TrPt sequential consortia experiment, h) the temperature and pH profiles throughout the operation of the SynCONS. *P. thermoglucosidasius* was inoculated after 7 days, i) the total secreted protein in the bioreactors, as indicated by the increase over time. The proteins reach their peak within 24 hours of cultures. j) the graph shows the released sugars (g (glucose equivalent/L)) in the culture by the enzymatic breakdown of cellulose, k) the HPLC spectra shows the difference between the day 0 and the Day 12 of the bioprocessing. The ethanol and lactic acid were quantified by measuring area under the respective peaks and comparing them with standards’ curves. The presence of ethanol and lactic acid establishes the proof of concept for the TrPt sequential bioprocessing, l) the percent g/g fermentation products yield in the TrPt bacterial co-culture consortia, with respect to utilised cellulose. The graphs in c), e), and j) have error bars representing the standard error of mean (n=4 biological replicates).

### Bacterial-bacterial thermophilic SynCONS of *T. fusca* and *P. thermoglucosidasius* (TfPt) for cellulose to value-added compounds

Individually optimised parameters (Figure 3 and Suppl. Fig. 3) were used to grow *T. fusca* and *P. thermoglucosidasius* NCIMB 11955 bacterial-bacterial consortium for maximum sugar and fermentative product release in bioreactors. A 10 %(v/v) inoculum of *T. fusca* yielded the highest free sugars in culture (up-to 6 µM/L) and introduction pf *P. thermoglucosidasius* on the third day of the consortia bioprocess was found optimum for highest ethanol titres (16 mg/L) (Suppl. Fig. 3).

**Figure 3:**
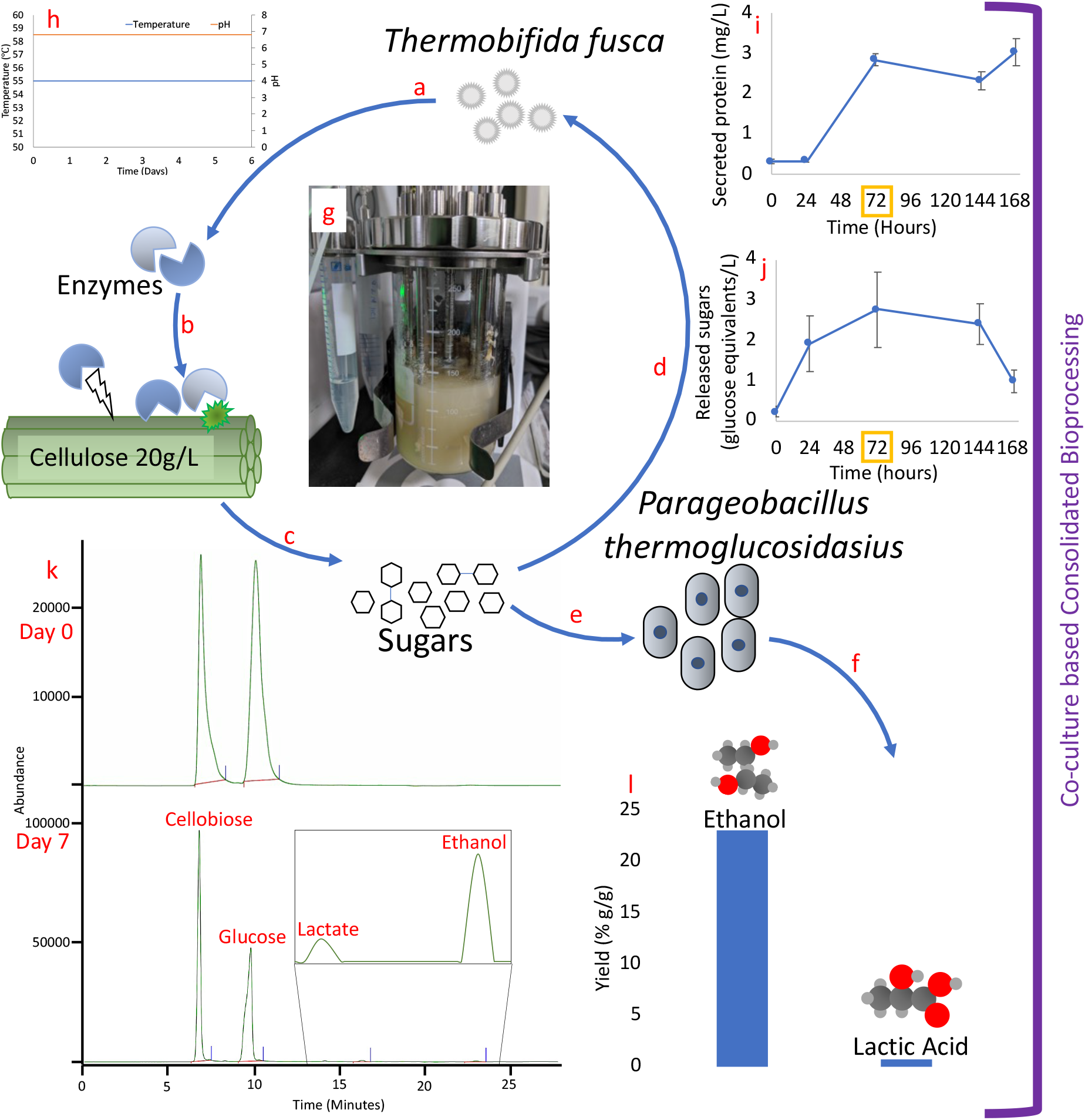
The bacterial-bacterial SynCONS of *Thermobifida fusca - Parageobacillus thermoglucosidasius* (TfPt) for cellulose to fermentative products. The co-culture consortium has both organisms co-existing and simultaneously performs saccharification of cellulose (by *T fusca)* and fermentation (by *P. thermoglucosidasius)* of the resulting monomeric and short oligomeric carbohydrates into lactic acid and ethanol. a) *T. fusca* produces and secretes the cellulases enzymes, b) the enzymes start to attack and break down the cellulose, c) the simpler sugars like glucose and cellobiose are released from cellulose breakdown, d) the sugars are taken up by *T. fusca* for growth, metabolism and enzyme production, e) Concurrently, *P. thermoglucosidasius* also takes up the sugar, f) the sugars are being utilised by *P. thermoglucosidasius* for the valuable chemicals production; ethanol and lactic acid in this experiment. The following supporting information is also supplied: g) the bioreactor containing the TfPt consortia co-culture experiment, h) the temperature and pH profiles throughout the operation, i) the total secreted protein in the bioreactors. *P. thermoglucosidasius* was inoculated after 48 hours (indicated by orange square), j) the graph shows the total released sugars (g (glucose equivalent)/L) in the culture by the enzymatic breakdown of cellulose, k) the HPLC spectra shows the difference between the day 0 and the day 7 of the bioprocessing. The ethanol and lactic acid were quantified by measuring area under the respective peaks and comparing them with standards’ curves. The presence of ethanol and lactic acid validates the co-culture consortia concept. l) The yield of fermentation products in the TfPt bacterial co-culture consortia, with respect to utilised cellulose (% g/g). The graphs in c), e) and j) have error bars representing the standard error of mean (n=4 biological replicates).

### Commercial *T. reesei* cellulase and *P. thermoglucosidasius* SSF of cellulose enhanced the lactic acid and ethanol yields

A comparison of commercial *T. reesei* cellulase at concentrations of 7, 14, 21 and 28 units/mL similar total sugars (mainly glucose and cellobiose) released at all concentrations. The optimised time for enzymatic saccharification was found to be 24 hours (Suppl. Fig. 8). We hypothesise that the low variation in sugar release could be a result of feedback inhibition of the enzymatic breakdown of cellulose by the reaction end products and intermediates like glucose and cellobiose respectively. As a result, an enzyme concentration of 7 unit/mL was used for cellulose degradation in subsequent experiments for it offered minimum costs associated with enzyme requirement.

The enzyme-bacterial bioprocessing combined the release of sugars in the bioreactor by the enzymatic breakdown of cellulose with fermentation of these sugars by *P. thermoglucosidasius* (Figure 4). The HPLC analysis showed the resultant titres of 0.19 g/L and 0.35 g/L for ethanol and lactic acid which correspond to 0.06 g/g ethanol and 0.12 g/g lactic acid yields respective to utilised cellulose.

**Figure 4:**
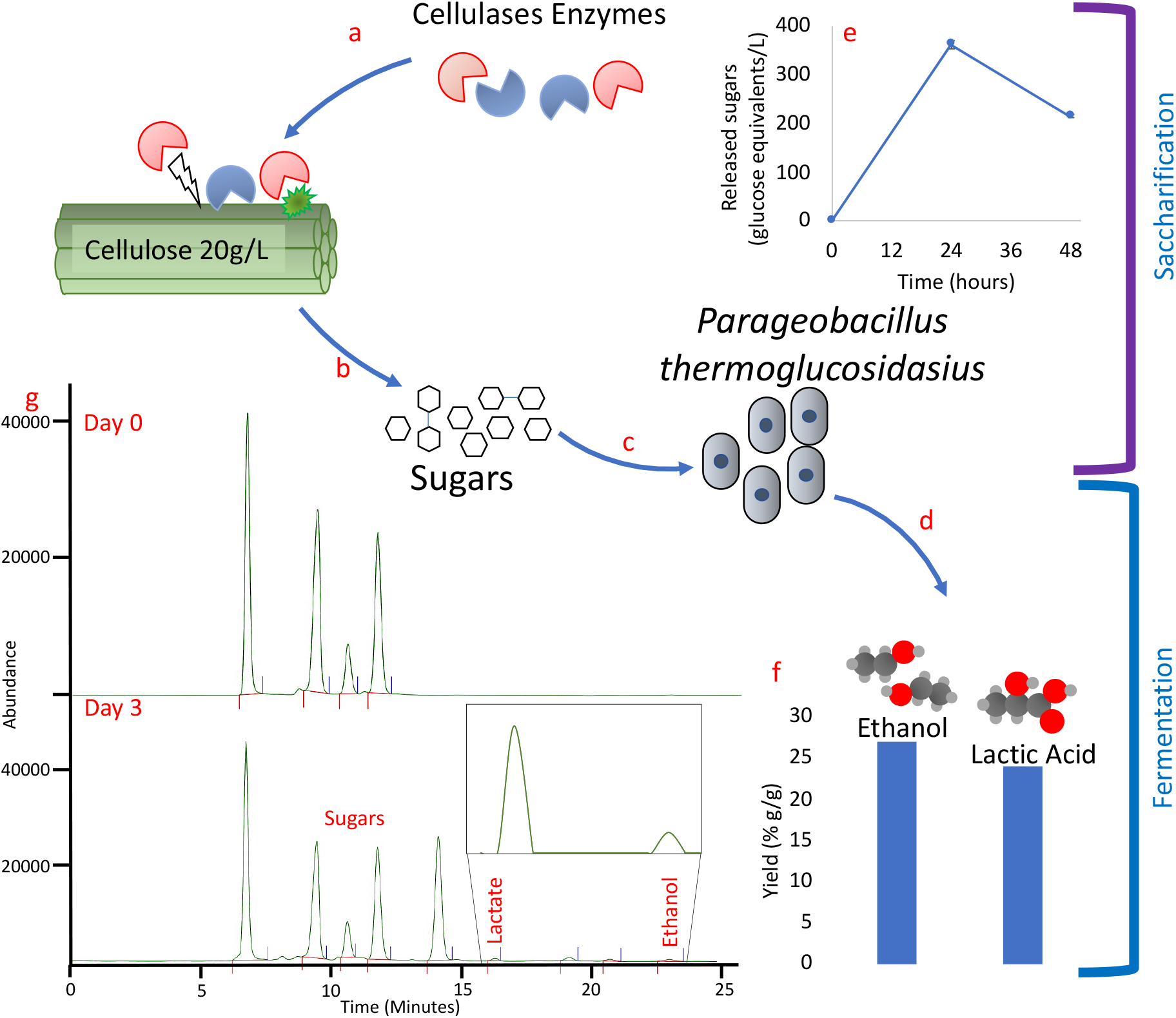
The commercial *T. reesei* cellulases and *P. thermoglucosidasius* bioprocess facilitates simultaneous saccharification and fermentation of the cellulose. a) Commercial *T. reesei* cellulases enzymes are added to the bioreactor, along with Avicel and ASM at 4.8 pH, b) the enzymes break down the cellulose into simpler sugars, c) after 24 hours, *P. thermoglucosidasius* is introduced, d) the sugars are being utilised by *P. thermoglucosidasius* for the production of target products ethanol, and lactic acid in this experiment. The following are the supporting data and results of different steps of the experiment: e) the graph shows the released sugars (glucose equivalent) in the culture by the enzymatic breakdown of cellulose, g) the HPLC spectra shows the difference between the day 0 and the Day 3 of the bioprocessing. The sugars at time 0 are released from autoclaving the culture media (containing avicel) f) Percent g/g yield of fermentation products in the enzyme-bacterial bioprocessing, with respect to utilised cellulose. (n=1).

### Pyrolysis of residual cellulose and microbial biomass post SynCONS bioprocessing resulted in valuable industrial carbon

Bioprocessing from the TrPt and TfPt SynCONSs resulted in 39% and 30% cellulose degradation respectively. Therefore, 61% and 70% of the cellulose remained unutilised, in addition to the resulting microbial biomass of the SynCONS bioprocess. This combined waste material was subjected to pyrolysis, which resulted in solid carbon with and overall weight loss of between 70 ± 10 % for dried samples.

Solid carbon material obtained by pyrolysis of the residual biomass (consisting of cells and unutilised cellulose) was characterised by SEM imaging, XRD and Raman spectroscopy (Figure 5). The pyrolyzed material retained its initial morphology, but with shrinkage in the dimensions (Figures 5a and 5b). The produced Raman spectrum (Figure 5 c) features the D-band (A_1g_ breathing mode of the hexagonal rings at the K-point, resulting from disorder) and the G-band (E_2g_ mode at the Γ-point, resulting from stretching vibrations in *sp*^2^ carbons) band at 1350 and 1560 cm^-1^ respectively. In addition, a small broad peak ranging from 2500-3000 cm^-1^ can also be observed, which is indicative of the presence of aromatic structures. Such Raman spectra are characteristic of most non-graphitizing polymeric carbon materials. ^72^ Natural polymeric precursors are known to produce porous carbon, which can be activated via physical or chemical routes.^38^

**Figure 5:**
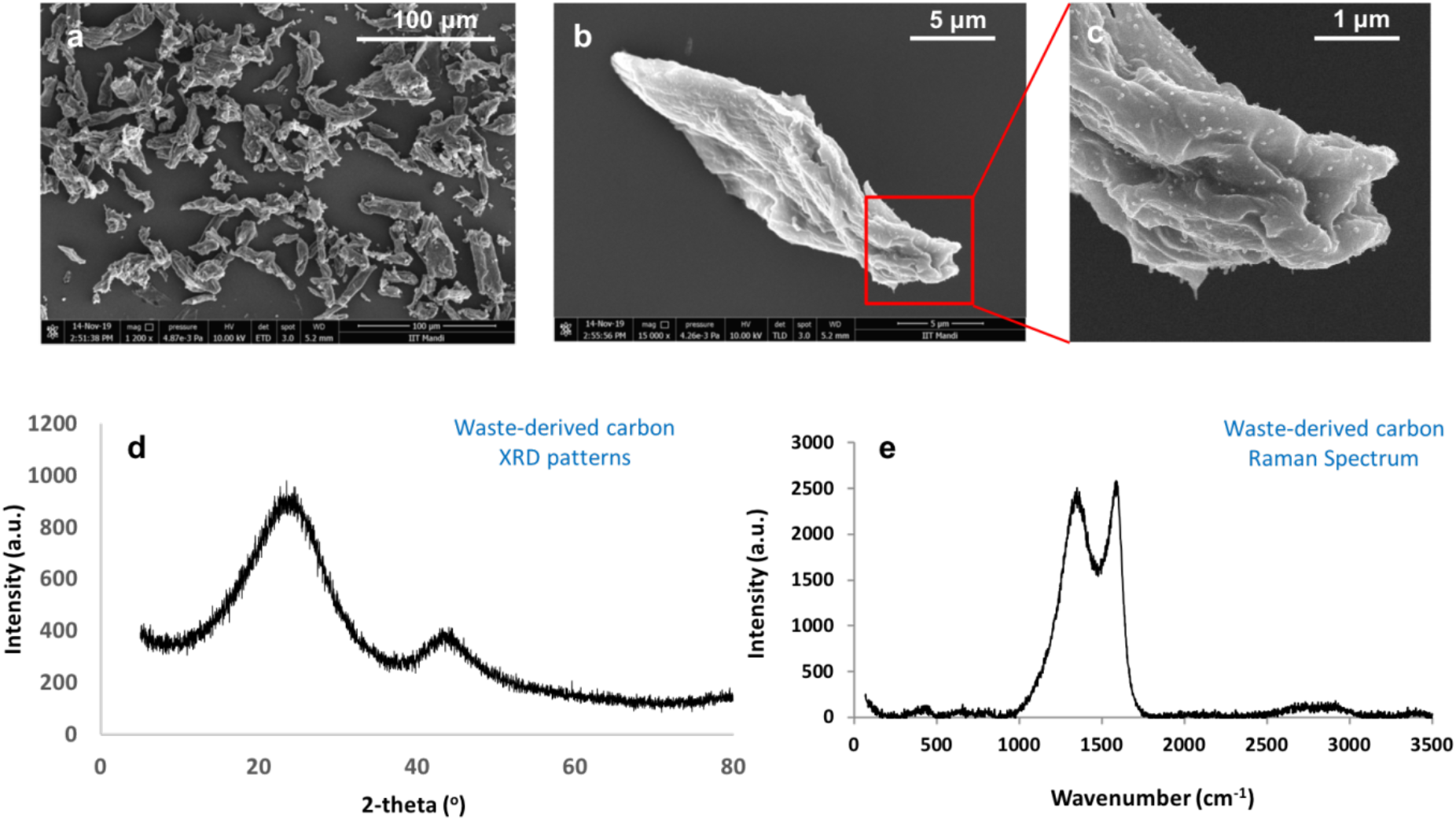
a)-c): SEM micrographs of pyrolyzed biomass at different magnifications; Scale bars: 100 µm, 5 µm, and 1 µm. d) XRD pattern; 2-theta is the measured angle between the incident and diffracted X-rays and e) Raman spectrum of carbon obtained after pyrolysis of solid residues from cellulose bioprocessing.

The XRD pattern of this carbon is shown in Figure 5 d). The broad peak ranging from 2θ values 10-35° is indicative of the (002) graphite basal planes. One can deduce that the material only features a short-range order. The second broad peak between 2θ values 40 and 50 corresponds to the (100) planes. Its Gaussian nature supports the idea of a highly disordered material with variable interlayer distances.

## Discussion

Hybrid technologies are a potentially useful approach for sustainable conversion of cellulose to valuable products. In nature, intricate microbial communities are formed to breakdown and utilise a complex polymer like cellulose. Synthetic consortia (SynCONS)-based consolidated bioprocessing has shown potential as an efficient approach to produce valuable chemicals from cellulosic feedstocks, where different microbes are specialised for different tasks.^3,5,7,12,15,73^ However, some of the unutilised cellulose remains, along with the microbial cells themselves used for the consolidated bioprocessing. This leftover material poses disposal challenges, especially when genetically modified organisms are involved in a large-scale bioprocess. In parallel, pyrolysis is another technique for extracting valuable compounds from cellulosic materials. Here, we have presented and validated a hybrid approach of integrating synthetic microbial consortia-based bioprocessing with pyrolysis for efficient valorisation of biomass. Fermentations were carried out for each SynCONS, producing lactic acid and ethanol, and the integration of pyrolysis of the residual carbon ensured complete use of the cellulose starting material in this complete bioprocess, mitigating the need of disposal of unutilised cellulose and cell biomass at the end of bioprocessing.

The SynCONS concept allows for a greater range for both bioprocess design and production of biomolecules, particularly when breakdown of complex molecules is involved. This has resulted in various studies employing a range of different approaches, including: same and multi species consortia, co-cultures and sequential consortia, multi-domain consortia, thermophile-mesophile consortia, and consortia with or without physical separation. However, the fermentation yields of SynCONS are typically found to be lower when compared to monocultures, primarily because of the multiple microbes involved both having their own ideal metabolic needs and competition for substrate. Theoretically, it may be possible to select any of the 218 short-listed organisms as consortium partners. The major challenging aspect of microbial co-cultures is fine-tuning the bioprocess to simultaneously support both saccharification and fermentation, which can have the added hindrance of maintaining sub-optimal growth parameters for individual consortia partners which enables their combined compatibility.^35^

In this study, aerobic organisms were prioritised as cellulose degrading SynCONS partners due to challenges in absolute anaerobic bioprocessing in a multi-phase consortium. An additional advantage of the aerobic degraders the production of extracellular free enzymes (as opposed to anaerobic microbes which mainly produce attached cellulosomes) as potential value-added products. ^74,75^ Two microbes (*Geobacillus genomosp. 3* and *Cellulosimicrobium cellulans* PSBB019) were identified from the leading organisms which has potential for both cellulose degradation and fermentation, which can be further tested as part of SynCONS or individual consolidated bioprocessing strains. However, the oxygen availability (aerobic/anaerobic conditions) during bioprocessing of cellulose would influence the metabolic priorities. Although, there are some reports of *Geobacillus* being cellulolytic, in practice they are poor at degrading crystalline cellulose (they mainly secrete endoglucanase). ^76–78^ While these microbes have great potential for CBP engineering, the main goal of this work was to design synthetic consortia with division of labour between specialised microbes for cellulose degradation and fermentation. The availability of these strains in microbial banks posed another challenge.

Based on these considerations, we designed a fungal-bacterial consortium and a bacterial co-culture consortium from the selected organisms. In the fungal-bacterial sequential consortium *T. reesei* produced cellulases required for cellulose break down and *P. thermoglucosidasius* fermented the resultant sugars into value-added products.

In the thermophilic bacterial co-culture consortium *T. fusca* was the cellulose degrader and *P. thermoglucosidasius* again used as the sugar fermenter. The co-culture consortium should simultaneously produce cellulases, break down the cellulose and ferment the resultant sugars into valuable products. Further, both optimized designed consortia were demonstrated to operate effectively in bioreactors, producing value-added products like lactic acid and ethanol.

Physicochemical parameters were the most influential constraints when designing these synthetic microbial consortia (SynCONS). The *T. reesei* and *P. thermoglucosidasius* TrPt consortia had to be used as a sequential consortium because of their differences in optimal growth conditions, leading to long bioprocessing times. However, the TfPt consortia could be used as a co-culture because of their similar growth conditions. Nevertheless, it was also necessary to fine tune the bioprocess parameters suitable for both saccharification and fermentation. The co-culture TfPt SynCONS allows microbial synergy between the partners, resulting in mutualistic benefits (Fig. 6). These mutualistic benefits were taken into consideration when designing the bioprocess of TfPt SynCONS. The excess oxygen in the bioreactors would be inhibitory to the fermentation, while the cello-oligosaccharides produced by the cellulose enzymatic breakdown are inhibitory to cellulase production and activity. In the TfPt SynCONS, *T. fusca* consumes the oxygen in the bioreactor, and the delayed introduction of *P. thermoglucosidasius* ensured a low dissolved oxygen throughout the remaining bioprocess, allowing fermentation. *P. thermoglucosidasius* was chosen as the fermentation partner for its metabolic abilities to consume glucose, cellobiose and larger cello-oligosaccharides alleviating feedback inhibition of cellulases production and activity (Daab and Leak, Unpublished). The bioprocessing parameters were also carefully selected to simultaneously allow cellulases production and fermentation (Suppl. Table 2).

**Figure 6:**
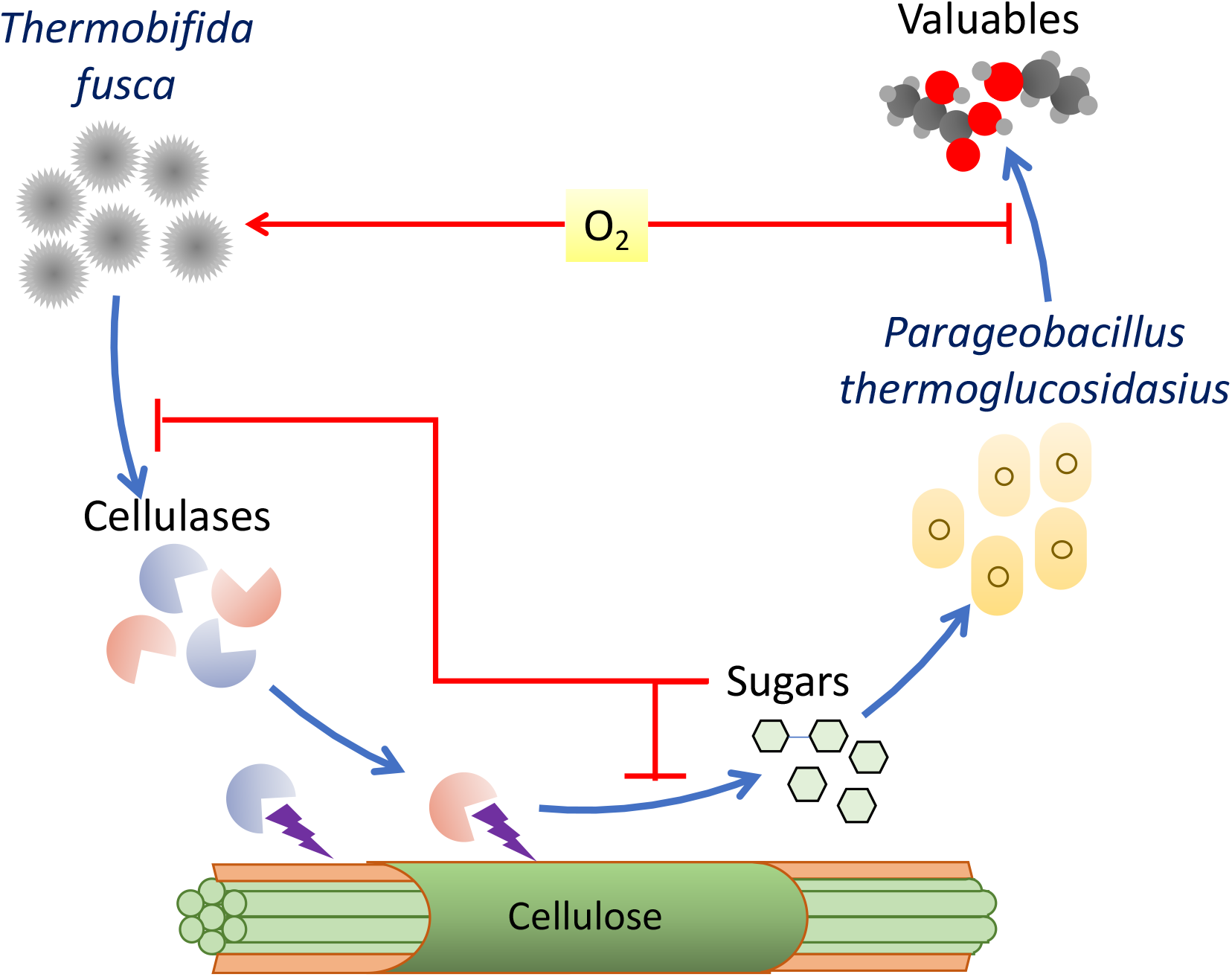
Synergetic action between *T. fusca* and *P. thermoglucosidasius* in coculture SynCONS. *T. fusca* helps *P. thermoglucosidasius* by consuming dissolved oxygen in the media, while *P. thermoglucosidasius* consume free sugars, including cello-oligosaccharides, thus reducing the inhibition of *T. fusca* cellulase production and activity.

Both the TrPt and TfPt consortia were compared to SSF (enzyme-bacterial bioprocessing) and lead to some interesting insights. Notably the microbial consortia consumed more cellulose compared to enzyme-bacterial bioprocessing, but the enzyme-bacterial consortia outperformed the microbial consortia in terms of target products titre. These results indicate that a significant proportion of the cellulose consumed by synthetic consortia is being utilised in cellular metabolism and maintenance of the consortium partners. While in the SSF, only the fermenting microbe consumes the resultant sugar, resulting in higher yields and titres. The cellulose breakdown profile of SSF differs significantly from the SynCONS, with the appearance of higher molecular weight oligosaccharides. We hypothesise the reasons are: a) use of commercial cellulase, and b) lack of cellulose degradation specialists (compared to SynCONS), which have the potential to consume higher molecular weight oligosaccharides. The higher yields of ethanol obtained by the TfPt242 consortium are expected from a specialised strain and further showcases the efficiency of designed consortia when specialised strains are used. (Suppl. Table 4)

*T. reesei* and *T. fusca*, the cellulose degraders employed in two SynCONS of this study have distinct cellulolytic systems (Suppl. Table 5), and cellulose breakdown mechanisms. While they both possess an array of cellulolytic enzymes, *T. reesei* is known to be limited by its β -glucosidase production.^79,80^ The *T. fusca* on the other hand, possesses unique processive cellulase TfCel9A, which mainly cleaves off cellotetraose from crystalline cellulose. TfCel48A is another unique exocellulase from *T. fusca*, which is proposed to work in synergy with TfCel9A, creating a positive feedback mechanism.^81,82^

The hybrid minimal-waste process involving high temperature pyrolysis of the spent fermentation culture, yielded a carbon material with physicochemical properties similar to commercial activated carbon.^83^ This could be employed for making water filtration columns, various adsorbents and even energy storage devices.^84^ As shown by the XRD and Raman analysis, the material features a short-range order and exhibits a low electrical conductivity, which can be further increased by increasing the pyrolysis temperature. The material occasionally also contained metal nanoparticles derived from the salts in the cell culture medium used. Such particles may induce specific functionalities in the carbon,^85^ which can be explored for sensor fabrication and other potential applications. The production of pyrolysis oils and syngas is relatively low when using an inert atmosphere; hence, their fractions were not isolated in this proof-of-concept study.

Although demonstrated in this study for bioprocessing of cellulose into ethanol and lactate, the SynCONS approach, could be applied to produce other industrial value-added chemicals. The research resulted in production of fermentation products from cellulose comparable to similar studies (Table 3), in addition to generating solid carbon for further commercial utilisation with minimal wastes in the process.

**Table 3:** Comparative assessment of various consortia based cellulose bioprocessing studies

| Consortia partners | Consortia type | Organism's type | Substrates used | Substrate treatment | Product(s) | Product titre | Product yield |
| --- | --- | --- | --- | --- | --- | --- | --- |
| *Trichoderma reesei* and *Parageobacillus thermoglucosidasius* (this study) | Fungal-Bacterial Mesophilic-thermophilic | Wild type | Avicel | No | Ethanol, Lactic Acid | 0.19 g/L and 0.05 g/L | 9.3 % g/g. |
| *Thermobifida fusca* and *Parageobacillus thermoglucosidasius* (this study) | Bacterial - Bacterial thermophilic | Wild type | Avicel | No | Ethanol, Lactic Acid | 0.36 g/L and 0.02 g/L | 23.7 % g/g |
| *Thermobifida fusca* and *Parageobacillus thermoglucosidasius* TM242(this study) | Bacterial - Bacterial thermophilic | Engineered | Avicel | No | Ethanol | 1.37 g/L | 32.7 % g/g |
| Commercial enzyme and *Parageobacillus thermoglucosidasius* (this study) | Enzyme-bacterial | Wild Type | Avicel | No | Ethanol, Lactic Acid | 0.19 g/L and 0.35 g/L | 51 % g/g |
| *T. reesei*/*E. coli*<sup>7</sup> | Fungal-Bacterial Mesophilic | Engineered | glucose, cellobiose, microcrystalline cellulose, or AFEX pre-treated corn stover | AFEX | Isobutanol | 1.88 g/L | 62 % |
| *Clostridium phytofermentans*/*Saccharomyces cerevisiae* and *Candida molischiana*<sup>16</sup> | Bacterial -Fungal Mesophilic | Engineered | α-cellulose | Addition of cellulases enzymes in fermentation | Ethanol | 22 g/L | 72 % (Ethanol + acetate + reducing sugars) |
| *Trichoderma reesei*, *Saccharomyces cerevisiae* and | Fungal-Fungal Mesophilic | Engineered | Avicel, Wheat straw | Acid hydrolysis in high pressure | Ethanol | 7.2 g/L | 67 % |
| *Scheffersomyces stipitis*<sup>3</sup> |  |  |  | autoclave at 140°C |  |  |  |
| *C. thermocellum* and *C. thermolacticum*<sup>86</sup> | Bacterial - Bacterial Thermophilic | Engineered | Glucose, xylose, cellobiose, microcrystalline cellulose, or hemicellulose | No | Ethanol | 4 g/L | 75 % |
| *Geobactersulfurreducens* and *Cellulomonas uda*<sup>87</sup> | Bacterial - Bacterial Mesophilic | Wild type | corn stover | AFEX | Electric potential, Ethanol | 1.9 mM Ethanol | 73 % of theoretical maximum |
| *Clostridium beijerinckii* and *Clostridium cellulovorans*<sup>88</sup> | Bacterial - Bacterial Mesophilic |  | alkali extracted deshelled corn cobs (AECC) | Alkali pretreatment | ABE | 2.64 g/L acetone, 8.30 g/L butanol and 0.87 g/L ethanol | 0.17 g/g (Calculated in this study) |
| *Clostridium thermocellum* and *Clostridium saccharoperbutylacetonicum* N1-4<sup>89</sup> | Bacterial - Bacterial Thermophilic-mesophilic | Wild type | Avicel | No | Butanol | 7.9 g/L | 40 % |
| *Clostridium* strain TCW1, *Bacillus* sp. THLA0409, *Klebsiella pneumoniae* THLB0409, *K. oxytoca* | “Thermophilic/thermotolerant” (60 °C) Mixed cultures | Wild type | Avicel and Napiergrass | acid-pretreatment method | Ethanol | NA | 0.108 g/g (Avicel) and 0.040 g/g |
| *THLC0409, and Brevibacillus strain AHPC8120*<br>90 |  |  |  |  |  |  | (Napier grass) |
| *Acremonium cellulolyticus C-1, and S. cerevisiae*<br>91 | Bacterial -Fungal | Engineered | Solka-Floc | No | Ethanol | 8.7-46.3 g/l | 0.15-0.18 g/g |

## Conclusions

The integrated process of SynCONS based microbial bioprocessing and pyrolysis demonstrated in this research was able to efficiently bioprocess cellulose into value-added products with minimal waste. We first established the designed SynCONS and sequential bioprocessing by producing lactic acid and ethanol from cellulose. Finally, subjecting the residual cellulose and microbial biomass to pyrolysis resulted in the production of carbon material with properties similar to commercial activated carbon. Future work in the field of synthetic consortia can use this workflow to optimise for other industrially relevant chemicals, especially those produced from biowaste.

## Acknowledgements

This work was supported by project BioPEC, jointly funded by the Department of Biotechnology (DBT), India (Ref. No. BT/IN/BMBF-Germany/29/SKM/2016-17) and the Federal Ministry of Education and Research (BMBF), Germany (Ref. No. 01DQ17014). CJ & JT thank the Ministry of Education, and IIT Mandi, India for doctoral research fellowship. CJ & JT are thankful to BioX centre and Advanced Materials Research Center, IIT Mandi for research facilities.

## Notes and references

‡ Footnotes relating to the main text should appear here. These might include comments relevant to but not central to the matter under discussion, limited experimental and spectral data, and crystallographic data.

§

§§

etc.

