## Supplementory Data File for "Integration of synthetic microbial consortia based bioprocessing with pyrolysis for efficient conversion of cellulose to valuables"

### 1. Supplementary figures

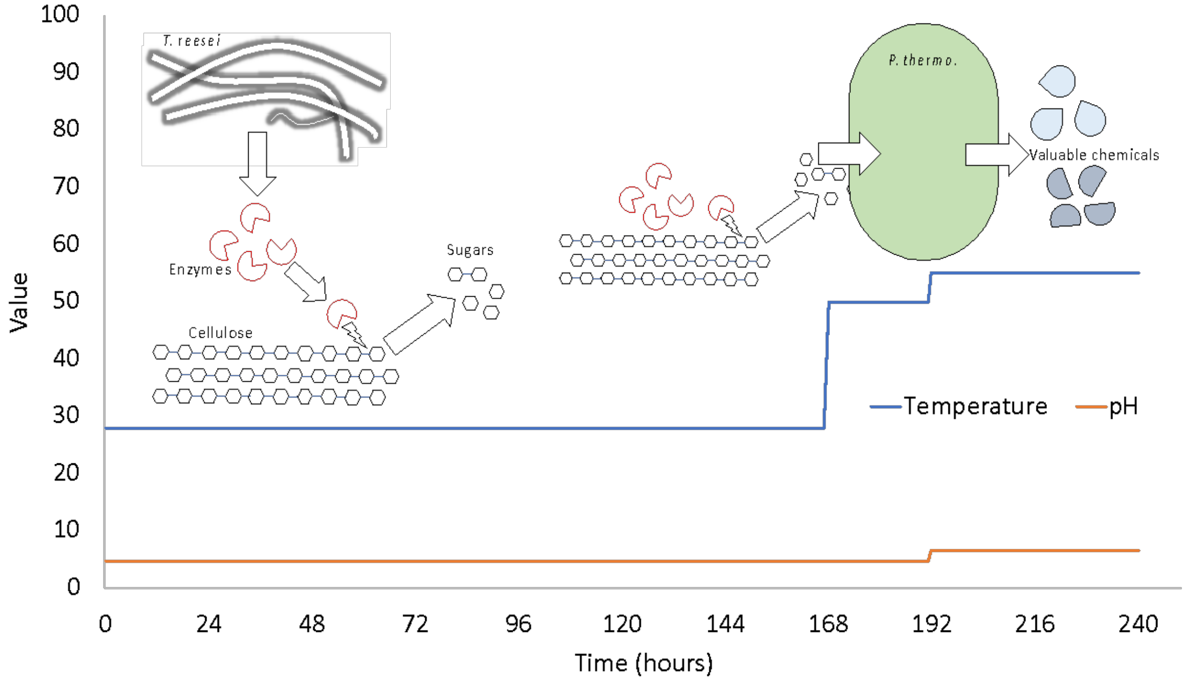


**Suppl. Fig. 1**: Bioprocessing parameters of TrPt fungal-bacterial sequential consortium in bioreactors. The changes in the pH and temperature in the saccharification and fermentation phases are reflected. At day 7, the temperature was increased to 50 °C from 28 °C for 24 hours. This will kill the fungus, but the fungal enzymes are optimum at 50 °C and will continue to work on the cellulose. After which the 2X ASM (without carbon source) media is added, temperature is set to 55°C and *P. thermoglucosidasius* is added.

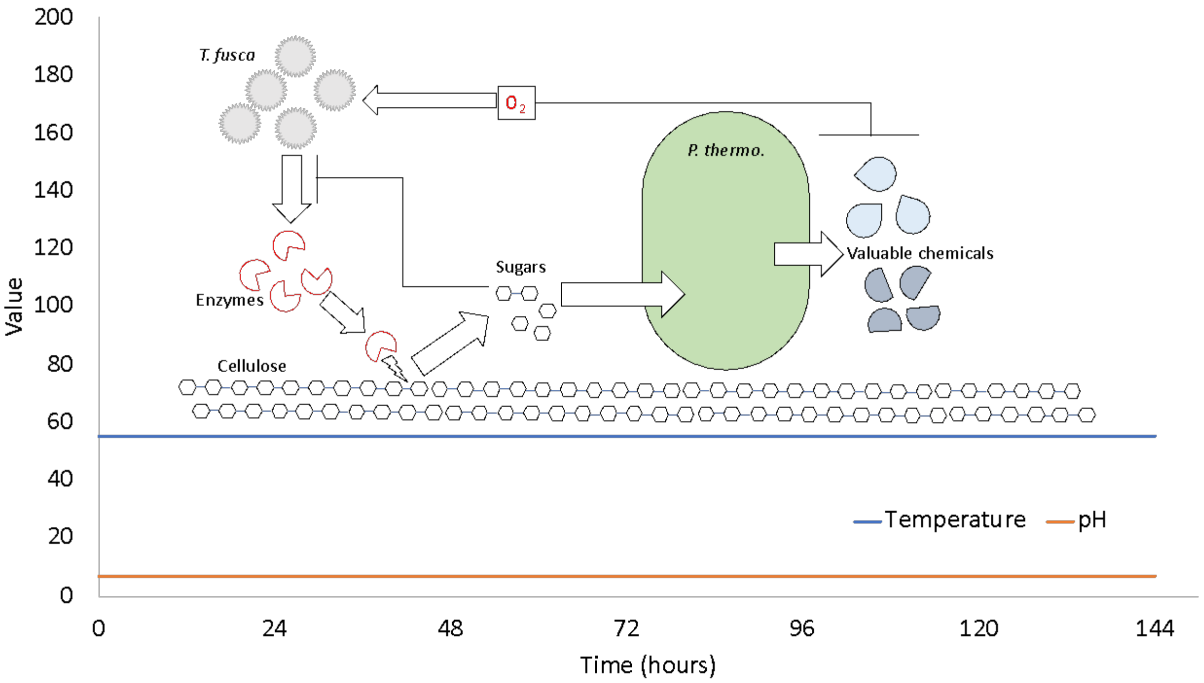


**Suppl. Fig. 2**: Bioprocessing parameters of bacterial-bacterial consortium in bioreactors. The growth parameters of *T. fusca* and *P. thermoglucosidasius* are very similar and maintained in the bioreactor throughout the bioprocess.

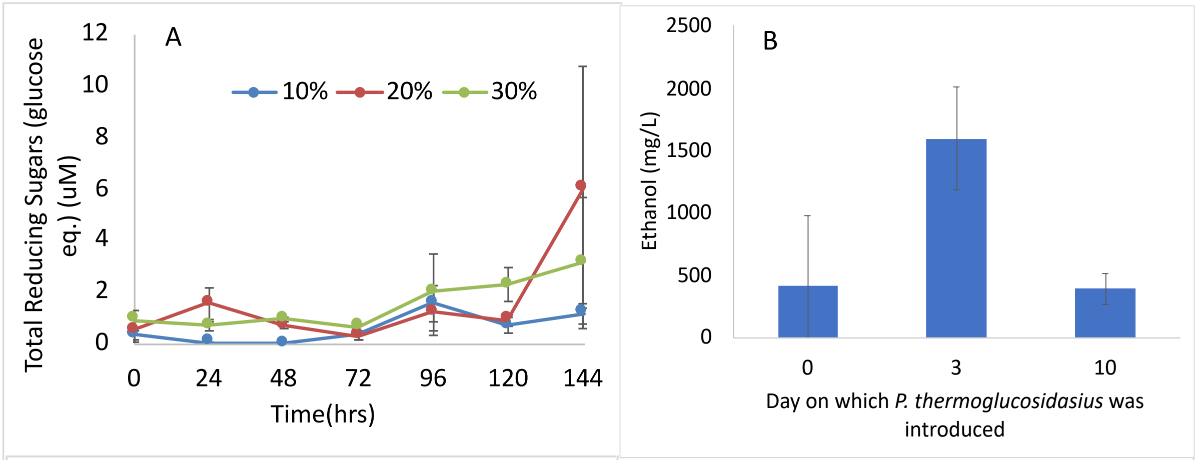


**Suppl. Fig. 3**: Optimisation of *T. fusca* inoculum. A. No significant difference was found in the released sugars with the increasing inoculum. So, 10% inoculum was used for the further bioreactor experiments. B. Optimisation of inoculation time of *P. thermoglucosidasius* in the consortia. Day 3 was fond to be suitable for the introduction of *P. thermoglucosidasius*.

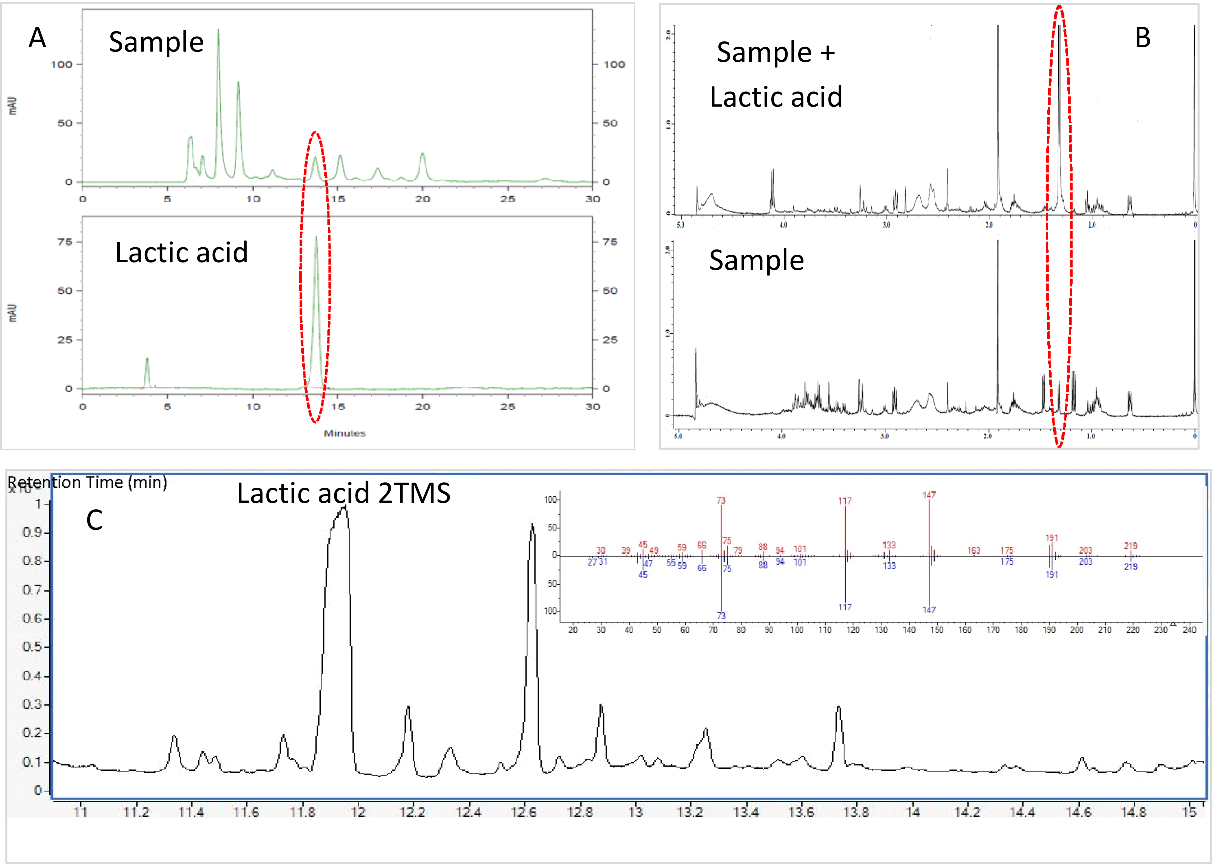


**Suppl. Fig. 4:** Identification of lactic acid from bioprocessed media through HPLC, GC-MS and H-NMR to confirm its production upon microbial bioprocessing of cellulose, (a) HPLC-chromatograms overlay of media sample after sequential microbial bioprocessing (microbial bioprocessing by *T. reesei* and *P. thermoglucosidasius*) of cellulose and lactic acid standard, (b) H-NMR spectral overlay of media sample after sequential microbial bioprocessing (microbial bioprocessing by *T. reesei* and *P. thermoglucosidasius*) of cellulose and its spiking with lactic acid standard, (c) GC-MS chromatogram of TMS derivatized media sample after sequential microbial bioprocessing (microbial bioprocessing by *T. reesei* and *P. thermoglucosidasius*) of cellulose

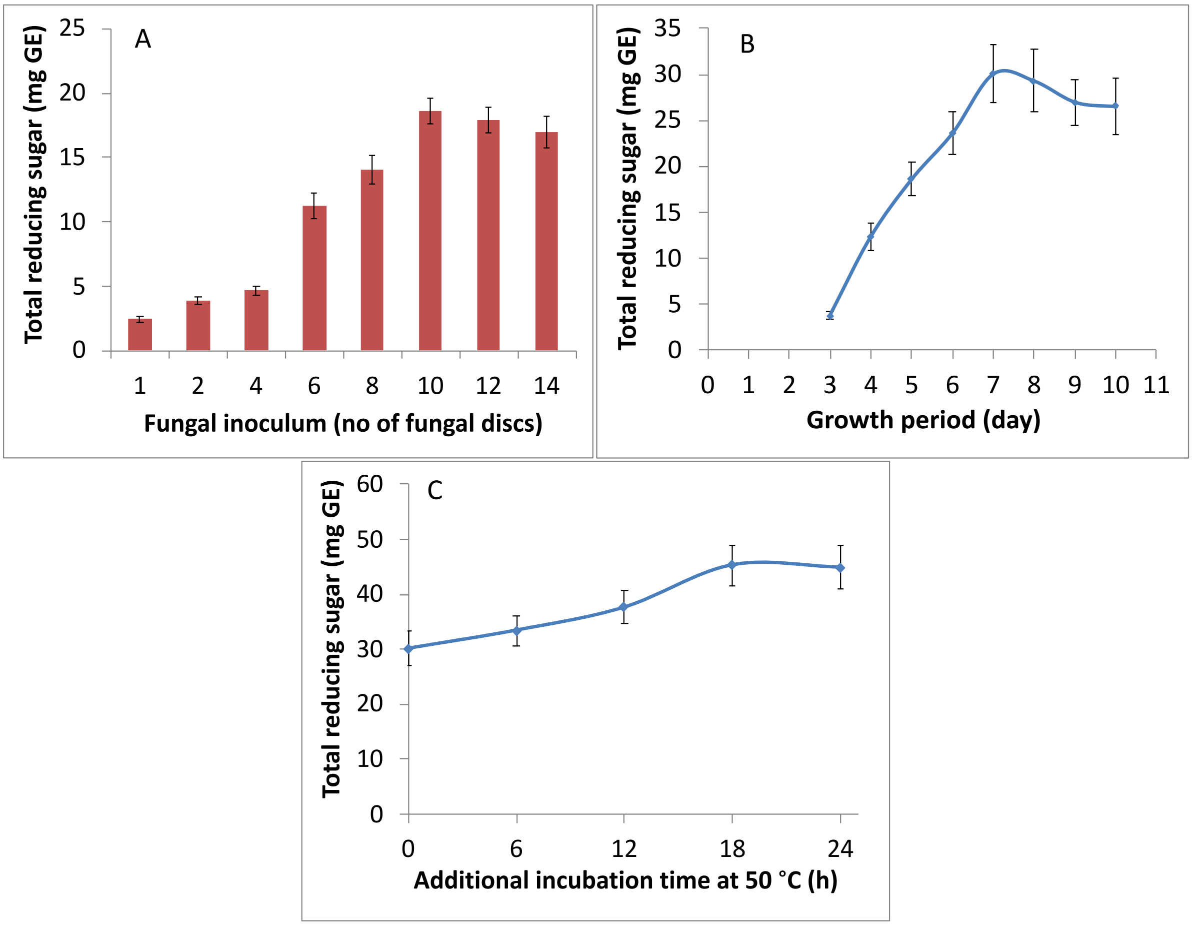


**Suppl. Fig. 5: A.** Optimization of fungal (*T. reesei* NCIM992) inoculum, B. growth/incubation time at 28 °C , and C. incubation time (time of incubation after the 7 day-growth of fungus) at 50 °C for maximum hydrolysis of cellulose into simple sugars. Hydrolysis of cellulose to simple sugar was measured in term of total reducing sugar content and each value is represented as mean ± SD (n = 4)

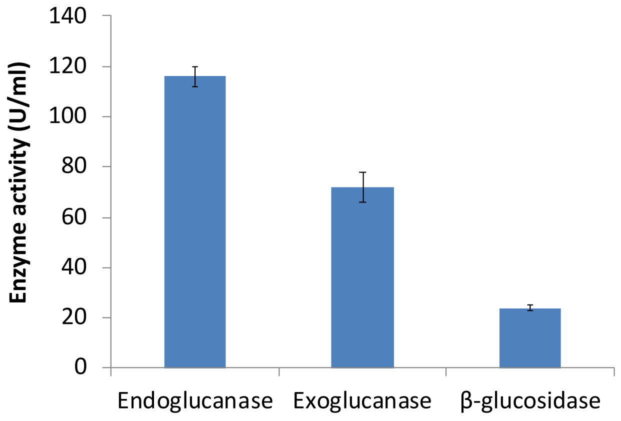


**Suppl. Fig. 6** Comparative account of endoglucanase, exoglucanase and *β*-glucosidase activity in media on 7th day of fungal (*T. reesei*) growth with initial inoculum 18.25 ± 2.21 mg (10 number of mycelial discs of 9.5 mm diameter), values are presented as mean ± SD (n = 4)

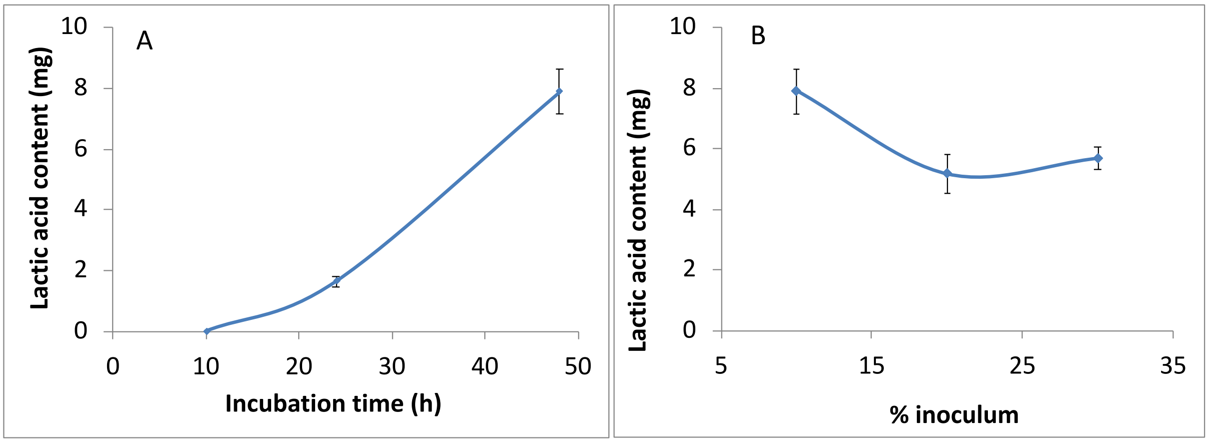


**Suppl. Fig. 7:** A. Optimization of *P. thermoglucosidasius* NCIMB11955 inoculum, and B. growth time for maximum utilization of glucose to produce lactic acid. HPLC was performed to quantify lactic acid and values are represented as mean ± SD (n = 4)

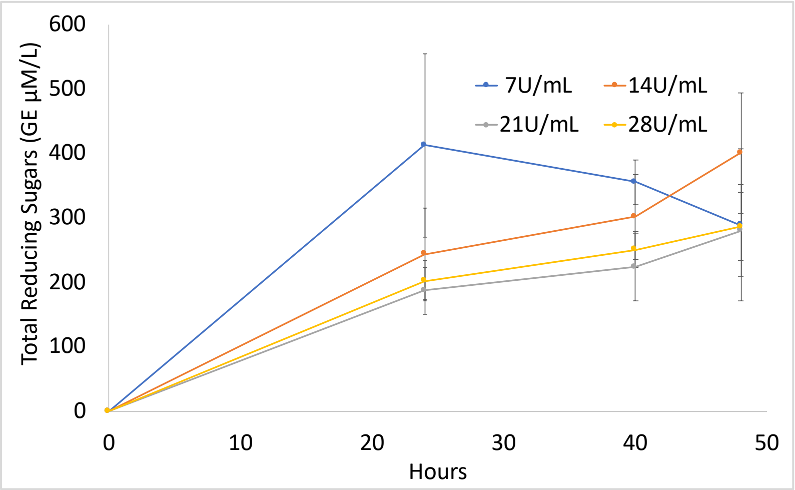


**Suppl. Fig. 8**: Optimisation of sugar release by commercial *T. reesei* enzymes. Increasing the enzyme units did not result in increased sugar released, so 7 units/mL enzymes concentration was selected for saccharification of cellulose for 24 hours.

**Suppl. Table S1**: Optimised and standardised bioprocessing parameters adopted for *T. reesei* and *P. thermoglucosidasius* SynCONS in bioreactors.

| Parameters | Saccharification phase | Fermentation phase (from 3rd day of experiment) |
|---|---|---|
| Media | 2x Hagerdahl Medium + 2% Avicel | 2x ASM (No carbon source), 90 mL |
| Total working volume | 100 mL | 180 mL |
| Temperature | 55 °C | 55 °C |
| pH | 7 | 7 |
| Dissolved Oxygen Tension | 20 % | 20 % |
| Air flow Actuator Limit | 1 VVM (100 mL/min) | 1 VVM for the first 4 hours (180 mL/Min)<br>0.05 VVM after 4 hours (9 mL/min) |
| Agitation | 200 RPM | 300 RPM |

**Suppl. Table S2**: Optimised and standardised bioprocessing parameters adopted for *T. fusca* and *P. thermoglucosidasius* SynCONS in bioreactors.

| Parameters | Saccharification phase | Fermentation phase (from 7th day of experiment) |
|---|---|---|
| Media | Trichoderma minimal medium + 2 % Avicel | 2x ASM (No carbon source), 90 mL |
| Total working volume | 100 mL | 180 mL |
| Temperature | 28 °C | 55 °C |
| pH | 4.8 | 7 |
| Dissolved Oxygen Tension | 20 % | 20 % |
| Air flow Actuator Limit | 1 VVM (100 mL/min) | 1 VVM for the first 4 hours (180 mL/Min)<br>0.05 VVM after 4 hours (9 mL/min) |
| Agitation | 200 RPM | 300 RPM |

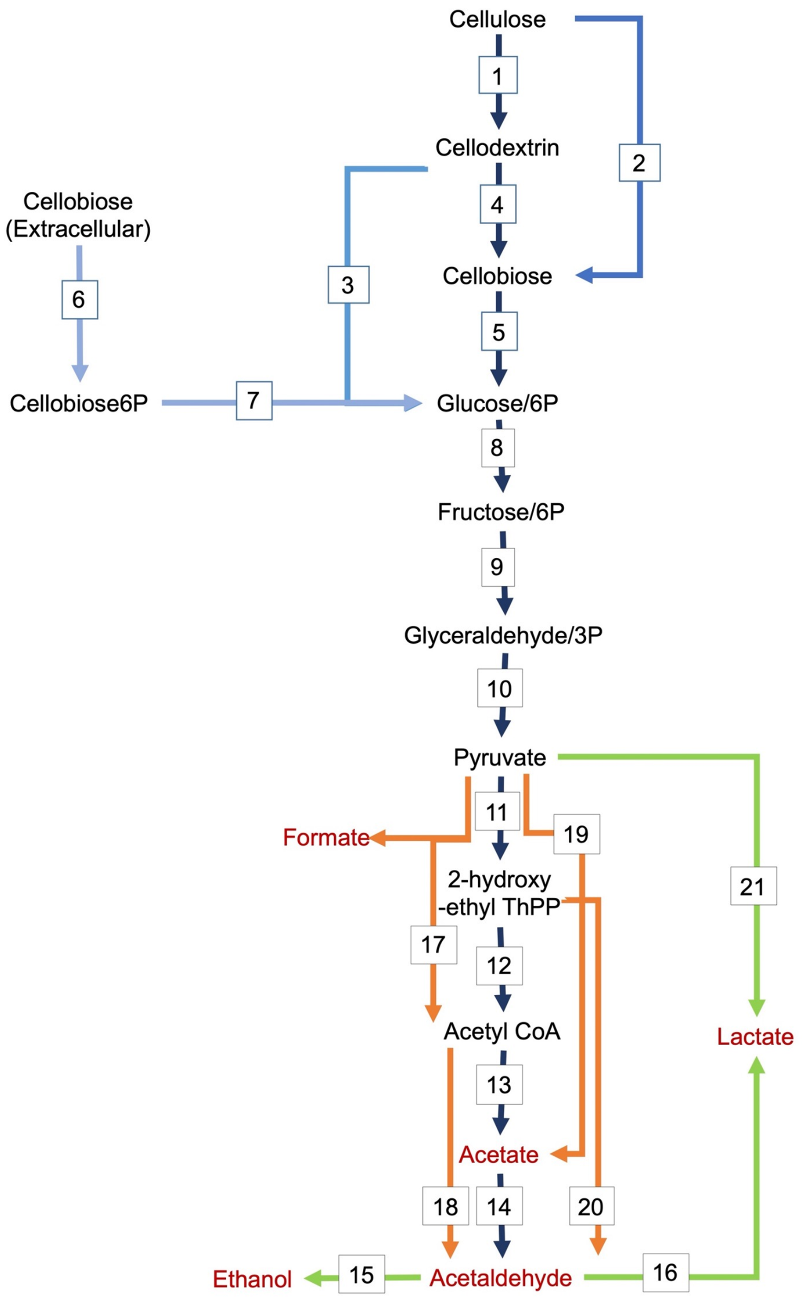


**Suppl. Figure 9:** Cellulose degradation and fermentation pathway. The reaction numbers and the presence and absence of the required genes for the reaction are matched with Suppl. Table S3.

**Suppl. Table S3: Comparison of cellulose degradation and fermentation metabolic capabilities of selected 218 organisms from KEGG database. Red indicates absence while the green indicates presence of enzyme required for the reaction.**

| KEGG code | Reaction Number (matched with Figure 4.4.1 A) → <br> Organism name and strain ↓ | 1 | 2 | 3 | 4 | 5 | 6 | 7 | 8 | 9 | 10 | 11 | 12 | 13 | 14 | 15 | 16 | 17 | 18 | 19 | 20 | 21 | Cellulose degradation completeness (%) | Fermentation completeness (%) |
|---|---|---|---|---|---|---|---|---|---|---|---|---|---|---|---|---|---|---|---|---|---|---|---|---|
| cth | *Hungateiclostridium thermocellum ATCC 27405* | | | | | | | | | | | | | | | | | | | | | | 57 | 57 |
| ctx | *Hungateiclostridium thermocellum DSM 1313* | | | | | | | | | | | | | | | | | | | | | | 57 | 57 |
| ace | *Acidothermus cellulolyticus* | | | | | | | | | | | | | | | | | | | | | | 71 | 50 |
| cac | *Clostridium acetobutylicum ATCC 824* | | | | | | | | | | | | | | | | | | | | | | 86 | 43 |
| cae | *Clostridium acetobutylicum DSM 1731* | | | | | | | | | | | | | | | | | | | | | | 86 | 43 |
| cay | *Clostridium acetobutylicum EA 2018* | | | | | | | | | | | | | | | | | | | | | | 86 | 43 |
| cpe | *Clostridium perfringens 13* | | | | | | | | | | | | | | | | | | | | | | 29 | 57 |
| cpf | *Clostridium perfringens ATCC 13124* | | | | | | | | | | | | | | | | | | | | | | 29 | 57 |
| cpr | *Clostridium perfringens SM101* | | | | | | | | | | | | | | | | | | | | | | 29 | 50 |
| ctc | *Clostridium tetani E88* | | | | | | | | | | | | | | | | | | | | | | 0 | 64 |
| ctet | *Clostridium tetani 12124569* | | | | | | | | | | | | | | | | | | | | | | 0 | 64 |
| cno | *Clostridium novyi* | | | | | | | | | | | | | | | | | | | | | | 0 | 50 |
| cbo | *Clostridium botulinum A ATCC 3502* | | | | | | | | | | | | | | | | | | | | | | 29 | 57 |
| cba | *Clostridium botulinum A ATCC 19397* | | | | | | | | | | | | | | | | | | | | | | 29 | 57 |

| | | | | | | | | | | | | | | | | | | | | | | | | |
|---|---|---|---|---|---|---|---|---|---|---|---|---|---|---|---|---|---|---|---|---|---|---|---|---|
| cbh | *Clostridium botulinum A Hall* | | | | | | | | | | | | | | | | | | | | | | 29 | 57 |
| cby | *Clostridium botulinum A2* | | | | | | | | | | | | | | | | | | | | | | 29 | 57 |
| cbl | *Clostridium botulinum A3 Loch Maree* | | | | | | | | | | | | | | | | | | | | | | 29 | 64 |
| cbk | *Clostridium botulinum B Eklund 17B (NRP)* | | | | | | | | | | | | | | | | | | | | | | 29 | 71 |
| cbb | *Clostridium botulinum B1 Okra* | | | | | | | | | | | | | | | | | | | | | | 29 | 64 |
| cbi | *Clostridium botulinum Ba4* | | | | | | | | | | | | | | | | | | | | | | 29 | 64 |
| cbn | *Clostridium botulinum BKT015925* | | | | | | | | | | | | | | | | | | | | | | 14 | 57 |
| cbt | *Clostridium botulinum E3* | | | | | | | | | | | | | | | | | | | | | | 29 | 71 |
| cbf | *Clostridium botulinum F Langeland* | | | | | | | | | | | | | | | | | | | | | | 29 | 64 |
| cbm | *Clostridium botulinum F 230613* | | | | | | | | | | | | | | | | | | | | | | 29 | 64 |
| cbj | *Clostridium botulinum H04402 065* | | | | | | | | | | | | | | | | | | | | | | 29 | 64 |
| cbe | *Clostridium beijerinckii NCIMB 8052* | | | | | | | | | | | | | | | | | | | | | | 57 | 71 |
| cbz | *Clostridium beijerinckii ATCC 35702* | | | | | | | | | | | | | | | | | | | | | | 57 | 71 |
| cbei | *Clostridium beijerinckii NCIMB 14988* | | | | | | | | | | | | | | | | | | | | | | 57 | 71 |
| ckl | *Clostridium kluyveri DSM 555* | | | | | | | | | | | | | | | | | | | | | | 0 | 71 |
| ckr | *Clostridium kluyveri NBRC 12016* | | | | | | | | | | | | | | | | | | | | | | 0 | 71 |
| clj | *Clostridium ljungdahlii* | | | | | | | | | | | | | | | | | | | | | | 0 | 64 |
| cls | *Clostridium sp. SY8519* | | | | | | | | | | | | | | | | | | | | | | 0 | 64 |
| clb | *Clostridium sp. BNL1100* | | | | | | | | | | | | | | | | | | | | | | 86 | 57 |
| csr | *Clostridium saccharoperbutylacetonicum* | | | | | | | | | | | | | | | | | | | | | | 86 | 64 |
| cpas | *Clostridium pasteurianum BC1* | | | | | | | | | | | | | | | | | | | | | | 29 | 57 |

| | | | |
|---|---|---|---|
| cpat | *Clostridium pasteurianum DSM 525 = ATCC 6013* | 14 | 64 |
| cpae | *Clostridium pasteurianum DSM 525 = ATCC 6013* | 14 | 64 |
| csb | *Clostridium saccharobutylicum* | 86 | 64 |
| cah | *Clostridium autoethanogenum* | 0 | 43 |
| clt | *Clostridium bornimense* | 57 | 57 |
| cbv | *Clostridium baratii* | 29 | 43 |
| csq | *Clostridium scatologenes* | 57 | 57 |
| cld | *Clostridium sporogenes* | 29 | 57 |
| cace | *Clostridium aceticum* | 0 | 50 |
| cck | *Clostridium carboxidivorans* | 71 | 57 |
| cbut | *Clostridium butyricum* | 29 | 64 |
| ctyk | *Clostridium tyrobutyricum* | 0 | 43 |
| ceu | *Clostridium estertheticum* | 71 | 64 |
| ctae | *Clostridium taeniosporum* | 29 | 43 |
| cfm | *Clostridium formicaceticum* | 0 | 50 |
| cchv | *Clostridium chauvoei* | 29 | 57 |
| carg | *Clostridium argentinense* | 29 | 64 |
| cdrk | *Clostridium drakei* | 29 | 57 |
| cia | *Clostridium isatidis* | 29 | 50 |
| csep | *Clostridium septicum* | 29 | 64 |
| cdy | *Clostridium diolis* | 57 | 64 |

| | | | |
|---|---|---|---|
| ccoh | *Clostridium cochlearium* | 0 | 36 |
| cfer | *Clostridium fermenticellae* | 0 | 50 |
| cgas | *Clostridium gasigenes* | 57 | 57 |
| ccel | *Clostridium cellulosi* | 57 | 50 |
| kll | *Klebsiella sp. LTGPAF-6F* | 57 | 86 |
| klw | *Klebsiella huaxiensis* | 57 | 86 |
| kok | *Klebsiella aerogenes KCTC 2190* | 57 | 86 |
| koc | *Klebsiella aerogenes EA1509E* | 57 | 86 |
| bbe | *Brevibacillus brevis* | 29 | 79 |
| blr | *Brevibacillus laterosporus* | 29 | 79 |
| bfm | *Brevibacillus formosus* | 29 | 79 |
| bagr | *Brevibacillus agri* | 29 | 79 |
| brw | *Brevibacillus sp. 7WMA2* | 29 | 79 |
| bchs | *Brevibacillus choshinensis* | 29 | 71 |
| gsu | *Geobacter sulfurreducens PCA* | 0 | 43 |
| gsk | *Geobacter sulfurreducens KN400* | 0 | 50 |
| cfl | *Cellulomonas flavigena* | 71 | 79 |
| cfi | *Cellulomonas fimi* | 71 | 79 |
| cga | *Cellulomonas gilvus* | 71 | 86 |
| cez | *Cellulomonas sp. PSBB021* | 71 | 79 |
| celz | *Cellulomonas shaoxiangyii* | 71 | 79 |

| | | | |
|---|---|---|---|
| cej | *Cellulomonas sp. JZ18* | 71 | 79 |
| celh | *Cellulomonas sp. H30R-01* | 71 | 86 |
| hsc | *Acetivibrio saccincola* | 57 | 64 |
| ccl | *Hungateiclostridium clariflavum* | 57 | 64 |
| rch | *Ruminococcus champanellensis* | 57 | 57 |
| ral | *Ruminococcus albus* | 57 | 50 |
| afu | *Archaeoglobus fulgidus DSM 4304* | 0 | 29 |
| afg | *Archaeoglobus fulgidus DSM 8774* | 0 | 29 |
| pru | *Prevotella ruminicola* | 57 | 36 |
| pmz | *Prevotella melaninogenica* | 57 | 36 |
| pdn | *Prevotella denticola* | 57 | 36 |
| pit | *Prevotella intermedia* | 0 | 29 |
| pdt | *Prevotella dentalis* | 57 | 36 |
| pro | *Prevotella sp. oral taxon 299* | 0 | 36 |
| pfus | *Prevotella fusca* | 57 | 36 |
| peo | *Prevotella enoeca* | 29 | 36 |
| pje | *Prevotella jejuni* | 0 | 36 |
| poc | *Prevotella oris* | 57 | 36 |
| bcel | *Bacteroides cellulosilyticus* | 57 | 36 |
| bth | *Bacteroides thetaiotaomicron VPI-5482* | 29 | 50 |
| btho | *Bacteroides thetaiotaomicron 7330* | 29 | 43 |

| | | | |
|---|---|---|---|
| bfr | *Bacteroides fragilis YCH46* | 29 | 43 |
| bfs | *Bacteroides fragilis NCTC 9343* | 29 | 43 |
| bfg | *Bacteroides fragilis 638R* | 29 | 43 |
| bfb | *Bacteroides fragilis BOB25* | 29 | 50 |
| bhl | *Bacteroides helcogenes* | 57 | 36 |
| bxy | *Bacteroides xylanisolvens* | 57 | 43 |
| boa | *Bacteroides ovatus* | 57 | 43 |
| bcac | *Bacteroides caccae* | 29 | 43 |
| bcae | *Bacteroides caecimuris* | 29 | 36 |
| bzg | *Bacteroides zoogleoformans* | 57 | 36 |
| bhf | *Bacteroides heparinolyticus* | 57 | 36 |
| bis | *Bacteroides intestinalis* | 57 | 36 |
| bun | *Bacteroides uniformis* | 57 | 36 |
| bvu | *Bacteroides vulgatus* | 29 | 43 |
| pco | *Phanerochaete carnosa* | 71 | 64 |
| ate | *Caldicellulosiruptor bescii* | 57 | 43 |
| cob | *Caldicellulosiruptor obsidiansis* | 57 | 43 |
| chd | *Caldicellulosiruptor hydrothermalis* | 57 | 43 |
| cow | *Caldicellulosiruptor owensensis* | 57 | 43 |
| cki | *Caldicellulosiruptor kristjanssonii* | 57 | 43 |
| ckn | *Caldicellulosiruptor kronotskyensis* | 57 | 43 |

| | | | |
|---|---|---|---|
| clc | *Caldicellulosiruptor lactoaceticus* | 57 | 43 |
| ccha | *Caldicellulosiruptor changbaiensis* | 57 | 43 |
| ani | *Aspergillus nidulans* | 71 | 71 |
| afm | *Aspergillus fumigatus* | 71 | 71 |
| act | *Aspergillus clavatus* | 71 | 71 |
| nfi | *Aspergillus fischeri* | 71 | 64 |
| aor | *Aspergillus oryzae* | 71 | 64 |
| ang | *Aspergillus niger* | 71 | 71 |
| afv | *Aspergillus flavus* | 71 | 64 |
| fox | *Fusarium oxysporum* | 71 | 71 |
| ncr | *Neurospora crassa* | 71 | 71 |
| gte | *Geobacillus thermoleovorans CCB_US3_UF5* | 29 | 71 |
| gtk | *Geobacillus thermoleovorans KCTC 3570* | 29 | 64 |
| gtm | *Geobacillus thermocatenulatus* | 29 | 79 |
| gli | *Geobacillus lituanicus* | 29 | 71 |
| gtn | *Geobacillus thermodenitrificans* | 29 | 71 |
| gwc | *Geobacillus sp. WCH70* | 0 | 79 |
| gyc | *Geobacillus sp. Y412MC61* | 29 | 79 |
| gya | *Geobacillus sp. Y412MC52* | 29 | 79 |
| gct | *Geobacillus sp. C56-T3* | 29 | 79 |
| gmc | *Geobacillus sp. Y4.1MC1* | 29 | 79 |

| | | | |
|---|---|---|---|
| ggh | *Geobacillus sp. GHH01* | 57 | 79 |
| gjf | *Geobacillus genomosp. 3* | 86 | 79 |
| gea | *Geobacillus sp. 12AMOR1* | 14 | 71 |
| gel | *Geobacillus sp. LC300* | 14 | 64 |
| gse | *Geobacillus stearothermophilus* | 29 | 71 |
| gsr | *Geobacillus subterraneus* | 57 | 71 |
| gej | *Geobacillus sp. JS12* | 29 | 79 |
| gza | *Geobacillus zalihae* | 29 | 79 |
| ptl | *Parageobacillus thermoglucosidasius DSM 2542* | 14 | 79 |
| ptb | *Parageobacillus toebii NBRC 107807* | 0 | 79 |
| tmt | *Thermoanaerobacter mathranii* | 57 | 43 |
| tpd | *Thermoanaerobacter pseudethanolicus* | 57 | 43 |
| ttm | *Thermoanaerobacterium thermosaccharolyticum DSM 571* | 86 | 50 |
| tto | *Thermoanaerobacterium thermosaccharolyticum M0795* | 57 | 50 |
| txy | *Thermoanaerobacterium xylanolyticum* | 57 | 43 |
| tsh | *Thermoanaerobacterium saccharolyticum* | 86 | 43 |
| tbi | *Thermobispora bispora* | 71 | 64 |
| tcu | *Thermomonospora curvata* | 71 | 71 |
| dth | *Dictyoglomus thermophilum* | 71 | 36 |
| dtu | *Dictyoglomus turgidum* | 71 | 36 |
| aac | *Alicyclobacillus acidocaldarius subsp. acidocaldarius DSM 446* | 57 | 71 |

| | | | |
|---|---|---|---|
| aad | *Alicyclobacillus acidocaldarius subsp. acidocaldarius Tc-4-1* | 57 | 64 |
| tma | *Thermotoga maritima MSB8* | 71 | 43 |
| tmm | *Thermotoga maritima MSB8* | 71 | 43 |
| tmi | *Thermotoga maritima MSB8* | 71 | 43 |
| tmw | *Thermotoga maritima MSB8* | 71 | 43 |
| tmq | *Thermotoga maritima Tma100* | 71 | 43 |
| tmx | *Thermotoga maritima Tma200* | 71 | 43 |
| tpt | *Thermotoga petrophila* | 71 | 36 |
| trq | *Thermotoga sp. RQ2* | 71 | 36 |
| tnp | *Thermotoga naphthophila* | 71 | 36 |
| thq | *Thermotoga sp. 2812B* | 71 | 43 |
| thz | *Thermotoga sp. Cell2* | 57 | 43 |
| thr | *Thermotoga sp. RQ7* | 71 | 43 |
| sta | *Spirochaeta thermophila DSM 6192* | 57 | 43 |
| stq | *Spirochaeta thermophila DSM 6578* | 57 | 50 |
| mas | *Mahella australiensis* | 71 | 50 |
| cce | *Ruminiclostridium cellulolyticum* | 57 | 57 |
| cle | *Cellulosilyticum lentocellum* | 71 | 57 |
| cew | *Cellulosilyticum sp. WCF-2* | 71 | 57 |
| acel | *Anaerocolumna cellulosilytica* | 86 | 57 |
| xce | *Xylanimonas cellulosilytica* | 71 | 79 |

| | | | |
|---|---|---|---|
| xyl | *Xylanimonas allomyrinae* | 43 | 86 |
| cet | *Cellulosimicrobium sp. TH-20* | 43 | 86 |
| cceu | *Cellulosimicrobium cellulans* | 86 | 86 |
| scl | *Sorangium cellulosum So ce56* | 71 | 71 |
| scu | *Sorangium cellulosum So0157-2* | 71 | 64 |
| bco | *Bacillus cellulosilyticus* | 86 | 79 |
| ccau | *Chryseobacterium carnipullorum* | 29 | 64 |
| cben | *Chryseobacterium bernardetii* | 57 | 64 |
| cgle | *Chryseobacterium gleum* | 57 | 57 |
| ctai | *Chryseobacterium taihuense* | 57 | 57 |
| clac | *Chryseobacterium lactis* | 57 | 57 |
| ctak | *Chryseobacterium taklimakanense* | 29 | 50 |
| cjg | *Kaistella jeonii* | 29 | 57 |
| ccas | *Kaistella carnis* | 0 | 57 |
| cant | *Kaistella antarctica* | 29 | 57 |
| kda | *Kaistella daneshvariae* | 0 | 57 |
| eva | *Epilithonimonas vandammei* | 29 | 64 |
| ebv | *Empedobacter brevis* | 29 | 64 |
| bbl | *Blattabacterium sp. (Blattella germanica) Bge* | 0 | 21 |
| tsi | *Thermococcus sibiricus* | 29 | 43 |
| thv | *Thermococcus sp. 2319x1* | 57 | 43 |

| | | | |
|---|---|---|---|
| gka | *Geobacillus kaustophilus* | 29 | 57 |
| gth | *Parageobacillus thermoglucosidasius* | 29 | 71 |
| cthr | *Chaetomium thermophilum* | 71 | 64 |
| tfu | *Thermobifida fusca* | 86 | 50 |
| tre | *Trichoderma reesei* | 71 | 57 |
| ccb | *Clostridium cellulovorans* | 86 | 50 |
| csc | *Caldicellulosiruptor saccharolyticus* | 57 | 36 |
| pfu | *Pyrococcus furiosus* | 29 | 36 |
| pho | *Pyrococcus horikoshii* | 57 | 36 |
| tko | *Thermococcus kodakarensis* | 0 | 43 |
| sso | *Sulfolobus solfataricus* | 29 | 43 |
| tna | *Thermotoga neapolitana* | 71 | 36 |
| mtm | *Myceliophthora thermophila* | 71 | 64 |
| asc | *Acidilobus saccharovorans* | 29 | 14 |
| iag | *Ignisphaera aggregans* | 57 | 29 |

| **Suppl. Table S4**: Fermentation yields and titres of the designed consortia in this study. | | | | | | | | | | |
|---|---|---|---|---|---|---|---|---|---|---|
| | **Cellulose (Avicel) added (g/L)** | **Cellulose (Avicel) Utilised (g/L)** | **Cumulative Free Sugars in filtrate (µM GE/L)** | **Ethanol (g/L)** | **Lactate (g/L)** | **g/g yield with respect to utilised cellulose** | | **Percent g/g yield with respect to utilised cellulose** | | **Total yield** |
| | | | | | | **Ethanol** | **Lactate** | **Ethanol** | **Lactate** | |
| **Commercial enzymes from *T. reesei*** | 20 (123.5 mM/L) | - | 412 ± 143 in 24 hours (0.07 g/L) | - | - | - | - | - | - | |
| **Enzyme-Bac (n=1)** | 20 (123.5 mM/L) | 2.9 (18 mM/L) | 577 ± 16.8 in 2 days (0.10 g/L) | 0.19 ± 0.01 (4124 uM/L) | 0.35 ± 0.03 (3885 uM/L) | 0.07 | 0.12 | 27 | 24 | 51 |
| **Fungal-Bac** | 20 (123.5 mM/L) | 7.8 (48.1 mM/L) | 724.2 ± 29.3 in 7 days (0.13 g/L) | 0.19 ± 0.21 (4124 uM/L) | 0.05 ± 0.04 (555 um/L) | 0.02 | 0.00 | 8 | 1.3 | 9.3 |
| **Bac-bac** | 20 (123.5 mM/L) | 6.1 (38 mM/L) | 40.1 ± 0.01 in 3 days (0.007 g/L) | 0.36 ± 0.01 (7815 uM/L) | 0.02 ± 0.00 (222 uM/L) | 0.06 | 0.00 | 23 | 0.7 | 23.7 |

| **Suppl. Table S5:** Comparison of reported cellulases enzymes in *Trichoderma reesei* and *Thermobifida fusca.* | | |
|---|---|---|
| **Cellulase** | ***T. reesei*** [1] | ***T. fusca*** [2] |
| Cel1a | Yes | No |
| Cel1b | Yes | No |
| Cel5a | Yes | Yes |
| Cel5b | Yes | Yes |
| Cel6a | Yes | Yes |
| Cel6b | No | Yes |
| Cel7a | Yes | No |
| Cel7b | Yes | No |
| Cel9a | No | Yes |
| Cel9b | No | Yes |
| Cel12a | Yes | No |
| Cel48a | No | Yes |
| Cel61a | Yes | No |
| BgCl | Yes | Yes |

### 2. Media Composition

Potato Dextrose Agar medium was prepared as per manufacturer protocol *i.e.,* 39 grams/L. The Trichoderma Minimum Medium was prepared by mixing all its components except cellulose, thereafter pH 4.8 was obtained using KOH. Now, 10 ml media was distributed in each bioreactor tube of 50 ml capacity before adding the cellulose. Sigma Avicel (PH-101, 50µm particles) was used as a standard cellulose carbon source in required growth media. The components of TMM include cellulose (Avicel® PH-101) as carbon source, 20 (g/L); $NH_4SO_4$, 7.6 (g/L); $KH_2PO4$, 15 (g/L); $MgSO_4$, 2.4 mM; $CaCl_2$, 4.1 mM; $FeSO_4.7H_2O$, 0.005 (g/L); $MnSO_4.7H_2O$, 0.0016 (g/L); $ZnSO_4.7H_2O$, 0.0014 (g/L) and $CoCl_2$, 0.0037 (g/L). The 2TY liquid medium was prepared by adding tryptone (16 g/L), yeast extract (10 g/L) and NaCl (5 g/L) in DI water and pH was adjusted to 7. The composition of 2TY agar medium is the same as 2TY liquid medium and an extra component agar (1.4 %) added to it. The ASM is a minimal medium and it is composed of $NaH_2PO_4$ (20 mM), $K_2SO_4$ (10mM), citric acid monohydrate (8 mM), $MgSO_4$ (5 mM), $CaCl_2$ (0.08 mM), $Na_2MoO_4$. $2H_2O$ (1.65 µM), ammonium sulphate (25 mM), sulphate trace element stock solution (5 ml/L), biotin (12 µM) thiamine (12 µM) and glucose (0.5%) as carbon source. Sulphate trace element stock solution was separately prepared in DI water and its components consisted of concentrated $H_2SO_4$ (5 ml/L), $ZnSO_4.7H_2O$ (25 µM), $FeSO_4.H_2O$ (100 µM), $MnSO_4.H_2O$ (50 µM), $CuSO_4.5H_2O$ (5 µM), $CoSO_4.7H_2O$ (10 µM), $NiSO_4.6H_2O$ (16.85 µM) and $H_3BO_3$ (0.08 ml/L). The Hagerdahl medium used for *T. fusca* growth consisted of NaCl (1.5 g/L), (NH4)2SO4 (3.1 g/L), Na2HPO4 (9.1 g/L), KH2PO4 (0.9 g/L), EDTA (0.025 g/L), MgSO4.7H2O (0.1 g/L), ZnSO4.7H2O ( 0.004 g/L), FeSO4.7H2O (0.01 g/L), MnSO4.H2O (0.0076 g/L), CaCl2.2H2O (0.0132 g/L), Biotin (0.001 g/L), Thiamine (0.001 g/L) and Avicel (20g/L) as carbon source. The yeast-tryptone media and agar was used to maintain *T. fusca* in the lab and for primary growth and it consisted of Tryptone (3.0g/L), Yeast extract (3.0g/L), Gl
